# The genetic basis of the black pupae phenotype in tephritid fruit flies

**DOI:** 10.1101/2024.06.07.597636

**Authors:** Daniel F. Paulo, Thu N.M. Nguyen, Chris M. Ward, Renee L. Corpuz, Angela N. Kauwe, Pedro Rendon, Rocio E.Y. Ruano, Amanda A.S. Cardoso, Georgia Gouvi, Elisabeth Fung, Peter Crisp, Anzu Okada, Amanda Choo, Christian Stauffer, Kostas Bourtzis, Sheina B. Sim, Simon W. Baxter, Scott M. Geib

## Abstract

The remarkable diversity of insect pigmentation offers a captivating avenue for exploring evolution and genetics. In tephritid fruit flies, decoding the molecular pathways underlying pigmentation traits also plays a central role in applied entomology. Mutant phenotypes like the black pupae (bp) have long been used as a component of genetic sexing strains, allowing male-only release in tephritid sterile insect technique applications. However, the genetic basis of bp remains largely unknown. Here, we present independent evidence from classical and modern genetics showing that the bp phenotype in the GUA10 strain of the Mexican fruit fly, *Anastrepha ludens*, is caused by a large deletion at the *ebony* locus resulting in the removal of the entire protein-coding region of the gene. Targeted knockout of *ebony* induced analogous bp phenotypes across six tephritid species spanning over 50 million years of divergent evolution. This functionally validated our findings and allowed for a deeper investigation into the role of Ebony in pigmentation and development in these species. Our study offers fundamental knowledge for developing new sexing strains based on the bp marker and for future evolutionary developmental biology studies in tephritid fruit flies.

## Introduction

The color palette and patterns displayed by insects are perhaps the most striking evidence of their extraordinary diversity. Insect pigmentation has long fascinated biologists, leading to insights into their evolution, ecology, development, genetics, and physiology^1–3^. A deep understanding of molecular mechanisms underlying pigmentation pathways in insects has not only implications in evolutionary and developmental biology but also provides fertile grounds for the development of genetic-based methods in applied entomology. A textbook example of how mutations affecting pigmentation traits can be implemented in insect pest management comes from genetic studies in tephritid fruit flies.

With nearly 5,000 recognized species, the family Tephritidae (true fruit flies) is one of the largest groups within Diptera. In addition to its remarkable diversity and intriguing life-history traits, the family is renowned for accommodating some of the most invasive insects worldwide^4,5^. Owing to their generalist herbivorous (polyphagous) habits, a few species have become major horticultural and agricultural pests^6,7^. Females of these species lay their eggs inside a wide range of fruits and vegetables, where hatching larvae will feed to complete their development, resulting in significant consequences on commodity production and trade. These characteristics ensure tephritids a position in food security, sustainable agriculture, and conservation biology debates. The sterile insect technique (SIT), as a component of area-wide integrated pest management (AW-IPM) approaches, is among the most effective and environment-friendly tactics for controlling tephritid pests^8^. The technique relies on the continuous mass-release of irradiation-sterilized males into target areas to mate with wild females, leading to the production of infertile embryos and suppression of pest populations^9^. A key determinant to the success of SIT in tephritids has been the linkage of selectable traits to the male sex in genetic sexing strains (GSS). Puparium color based GSS have been developed in several fruit flies using the naturally occurring mutations *white pupae* (*wp*)^10–12^ and *black pupae* (*bp*)^13–15^. In these strains, females are homozygous for recessive mutations causing an atypical white or black puparium pigmentation, while males display the typical brown puparium color due to a chromosomal translocation of the wildtype rescue allele onto the male Y-chromosome, resulting in a consistent heterozygous state. This sex-linked color dimorphism allows female removal before releases, ensuring cost-effectiveness and efficacy of large-scale SIT programs^16^.

The white pupae (wp) phenotype in three distantly related tephritids, the Mediterranean fruit fly (medfly) *Ceratitis capitata*, the oriental fruit fly *Bactrocera dorsalis*, and the melon fly *Zeugodacus cucurbitae*, results from parallel mutations in a single, conserved gene encoding a Major Facilitator Superfamily (MSF) transporter protein^17^. This molecular identity supports early biochemical studies showing that the *wp* gene is essential to transport hemolymph catecholamines (pigment precursors) to the pupal cuticle^18^, promoting normal sclerotization and pigmentation. Despite gene conservation, similar *wp* mutations were never found in *Anastrepha*—a mega-diverse genus of fruit flies in the American (sub)tropics that includes major fruit-infesting pests, such as the Mexican fruit fly (mexfly) *Anastrepha ludens* and the South American fruit fly *Anastrepha fraterculus*. Alternatively, the black pupae (bp) mutant phenotype has been observed in *Anastrepha* and used to develop GSSs in this group of flies; however, its genetic basis remains largely unknown.

In this study, we present a collection of independent evidence from genetics, transcriptomics, and functional genomics, showing that *ebony* is the responsible gene for the mutant black pupae trait in *A*. *ludens* and likely other tephritids displaying parallel phenotypes. Our findings support the following conclusions: (1) the bp phenotype in the *A*. *ludens* GUA10 strain^13^ is caused by a large deletion at the *ebony* locus. This deletion removes the entire protein-coding region of the gene, thus disrupting its function; (2) disruption of *ebony* function is sufficient to recreate analogous bp phenotypes in diverse tephritids, indicating its potential as a candidate gene for other naturally occurring bp mutations within the family; (3) Ebony plays an essential role in inhibiting black melanization in adult fruit flies, which constitutes one of the genetic mechanisms underlying pigmentation differences within and between tephritid species; and (4) the *ebony* gene may have pleiotropic effects on both embryo viability and adult development in tephritids, ultimately impacting fitness. We discuss these discoveries through the lens of dipteran evolutionary developmental biology^19^ and their implications for the construction of new GSS for SIT applications.

## Results

### Generation of a mapping population

We adopted the *A*. *ludens* GUA10 strain as our model to investigate the genetic basis of the black pupae trait in tephritids (Fig. 1a). GUA10 is a GSS with an autosomal recessive *bp* mutation as a selectable marker, allowing sex separation based on pupal color dimorphism^14^. Females from GUA10 are homozygous for the *bp* mutation, exhibiting an atypical dark pupal case. The mutation also induces darkening of larva anal lobes and ectopic melanization of adult cuticle. Conversely, GUA10 males carry an irradiation-derived translocation between the Y-chromosome and the polytene chromosome 3 (mitotic chromosome 2), the latter known to harbor the bp-related gene^13^. This rearrangement links a functional *bp* allele to the male sex. Consequently, males are consistently heterozygous, displaying a wildtype brown pupal case.

**Fig. 1:**
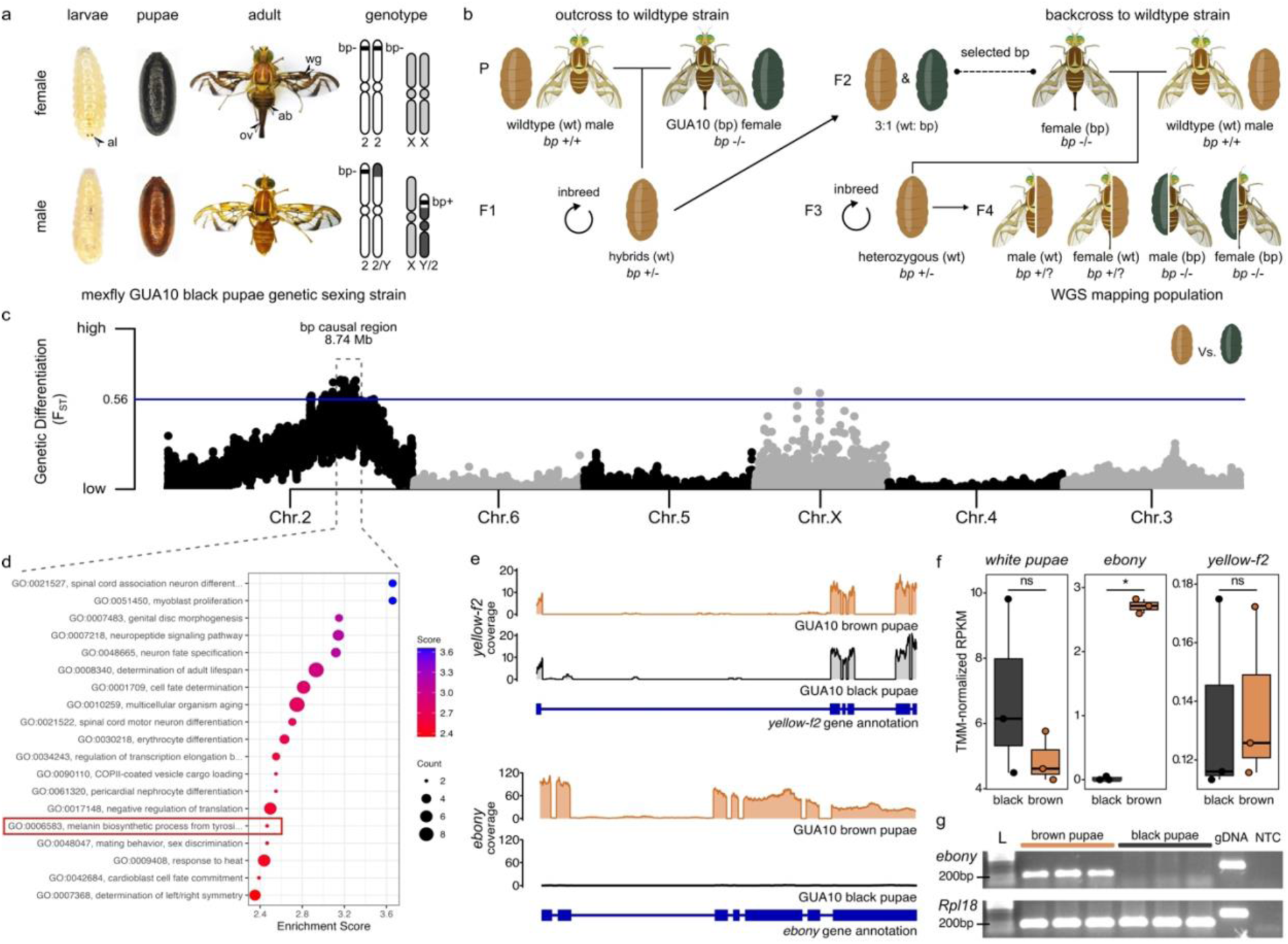
The *ebony* gene is linked to the black pupae phenotype in GUA10. **a** The bp phenotype is the sex-linked trait used for sex-sorting the mexfly GUA10 strain. Females from GUA10 exhibit black larval anal lobes (al) and pupal cases, developing into darker adults. Mutant adults are noticeable due to their darker wing stripes (wg), abdomen (ab), and ovipositor (ov). **b** To identify the loci responsible for the bp phenotype, we introgressed the GUA10 *bp* mutation (*bp*^−(GUA10)^) into a common wildtype genetic background. Subsequently, we established and whole-genome sequenced a F4 mapping population where siblings develop either wildtype brown (wt) or mutant black (bp) pupal cases. **c** Genetic differentiation (F_ST_) between black and brown pupae siblings (calculated for 100 Kb windows at 20 Kb sliding intervals across the reference genome) located a large interval within the *A*. *ludens* chromosome 2 showing significant differentiation. The horizontal line indicates the top 0.25th percentile threshold for significant intervals. **d** Enriched gene sets within this causal region include the genes *yellow-f2* and *ebony* from the melanin biosynthesis pathway; two promising candidates to explain the bp phenotype in GUA10. **e** RNA-Seq coverage revealed the silencing of *ebony* expression in black pupae females from GUA10. **f** The *ebony* gene is differentially expressed between black (homozygous females) and brown (heterozygous males) pupae siblings from GUA10, while no difference is detectable for *yellow-f2*. The expression of the *white pupae* gene, an MFS transporter required to provide pigment precursors to the pupal cuticle, is shown as a reference. Dots represent biologically independent replicates (*n* = 3), * = FDR < 0.05 and log2FC > 2, and ns = non-significant. **g** Semiquantitative RT-PCR assays further validated the RNA-Seq results. Amplifications of *Rpl18* are internal controls. L = 100 bp DNA ladder (NEB), gDNA = genomic DNA control, and NTC = non-template negative control.

We conducted a series of genetic crosses to introgress the *bp* mutation into a wildtype genetic background, enabling us to identify the relative genomic location of the bp causative loci in *A*. *ludens* using association mapping (Fig. 1b). We selected wildtype (wt) males to provide a reference background, and females from GUA10 as donors for the *bp* mutation (*bp*^−(GUA10)^). Isolated-mating between GUA10 females (*bp*^−/−^) and wt males (*bp*^+/+^) followed by inbreeding of F1 hybrids (*bp*^+/−^) led to the segregation of *bp* alleles at F2, placing them within a common genetic background. To further increase wildtype representation and recombination between divergent genomes, F2 bp females were individually backcrossed to wt males. Then, the resulting heterozygous F3 (*bp*^+/−^) individuals were inbred to generate the F4 mapping population, where siblings share similar genetic makeup with the exception of the *bp* mutation, and thus develop as black or brown pupae independently of their sexes.

### Association mapping implicates pigmentation genes to the black pupae phenotype in A. ludens GUA10

To map the *bp*^−(GUA10)^ mutation, we generated whole genome sequences (WGS) from black (*bp*^−/−^) and brown (*bp*^+/+^ or *bp*^+/−^) pupae siblings from the F4 mapping population (*n* = 18 of each phenotype). Resulting short reads were aligned to the *A*. *ludens* reference genome (GenBank: GCA_028408465.1) and used to identify DNA variants in each individual. We next calculated measures of genetic diversity along the genome using 100 kb sliding windows with a 20 kb step size. To minimize false identifications, we only considered the top 0.25% of windows from scans. Genome-wide landscape of pairwise genetic differentiation (F_ST_) between black and brown pupae siblings revealed a large island encompassing 8.74 Mb within the *A*. *ludens* chromosome 2 exhibiting the most significant genetic difference between phenotypes (Fig. 1c). Therefore, this genomic interval was considered the candidate region of the *bp*^−(GUA10)^ mutation.

To gain insight into the biological functions within this candidate region, we conducted a gene ontology (GO) enrichment analysis, comparing the gene content of its genomic interval to the complete genome annotation (Fig. 1d). Out of the 99 protein-coding genes located at the candidate region, seven exhibited significant enrichment in categories related to cuticle melanization and sclerotization (Supplementary Table 1). This set included five genes within the neuropeptide signaling pathway (Fisher’s exact test, *p* = 7.20e-4) and two genes involved in the melanin biosynthetic process from tyrosine (Fisher’s exact test, *p* = 3.42e-3). Upon closer examination, we identified *yellow-f2* (GenBank: XP_053946313.1) and *ebony* (GenBank: XP_053946246.1) as candidate genes associated with the black pupae phenotype in mexfly.

In *Drosophila melanogaster*, mutations in *yellow* result in a loss of black pigment, underscoring the essential role of the Yellow protein in black melanin production^20^. The *yellow-f2*, a member of the Yellow gene family, is implicated in melanization during later pupal and adult development^21^. In contrast, *ebony* mutants exhibit increased black pigmentation^22^. Previous studies have demonstrated that the pattern and intensity of melanization in *Drosophila* are largely determined by the coordinated expression of Yellow and Ebony^23^. Therefore, the melanin-promoting gene *yellow-f2* and the melanin-inhibiting gene *ebony* emerged as promising candidates responsible for the black pupae phenotype in *A*. *ludens*.

### ebony is silenced in black pupae females from A. ludens GUA10

We performed RNA-Seq transcriptome profiling on 1-day-old black (females, *bp*^−/−^) and brown (males, *bp*^+/−^) pupae siblings from the GUA10 strain currently maintained in Guatemala, under the assumption that transcriptional errors in candidate protein-coding genes result in the mutant trait. Similar to the approach used to identify the *wp* mutation^17^, we anticipated that differential read coverage between genotypes would reveal transcript variants or alterations in gene expression. RNA-Seq coverage was similar for *yellow-f2*; however, only reads from brown pupae males mapped to *ebony* (Fig. 1e). Differentially expressed gene (DEG) analysis showed significant downregulation of *ebony* in black pupae individuals (log_2_ fold-change = 7.4, *p* = 8.3e-06, and FDR = 0.03; Fig. 1f and Supplementary Table 2). Semiquantitative RT-PCR confirmed the silencing of *ebony* in black pupae females of GUA10 (Fig. 1g), leading to the hypothesis that *ebony* is the causative gene behind the mutant phenotype in this strain. Reinforcing this idea, *in situ* hybridization located *ebony* within region 24 of the polytene chromosome 3 (mitotic chromosome 2) in wildtype *A*. *ludens* (Supplementary Fig. 1), which is part of the 2-Y translocation chromosome found of both GUA10^14^ and its predecessor strain TAP-7^13^.

### The black pupae phenotype in A. ludens GUA10 results from a large indel variant at the ebony locus

We next screened the genomic regions surrounding *ebony* for DNA variants called from WGS short-reads of the F4 mapping population. Apart from single nucleotide and short insertion-deletion polymorphisms (SNPs and indels, respectively) in adjacent sequences, we could not identify variants within *ebony* (Fig. 2a), indicating that either this locus is identical between brown and black pupae siblings of the mapping population or there is a lack of read coverage in one or both subpopulation phenotypes. To better understand this pattern, we examined the short-read mapping coverage and identified a broad chromosomal interval with no coverage from black pupae individuals (Fig. 2b, Supplementary Fig. 2). This observation suggested that most *ebony* loci are missing in these flies. To confirm this putative deletion event, we sequenced a single male and female of *A*. *ludens* GUA10 strain with PacBio HiFi sequencing. We reasoned the effort could provide evidence of the *bp*^−(GUA10)^ mutation in single long HiFi reads associated with *ebony*. Both male and female samples generated approximately 69 Gb of data, representing over 80× coverage of the estimated 820 Mb genome. Mapping of HiFi reads to the reference genome of *A*. *ludens* validated our assumption and revealed a complex DNA variant spanning a 20,182 bp region within the *ebony* locus (Fig. 2c, Supplementary Fig. 2). The variant contains two large deletions of 8,186 bp and 4,310 bp, resulting in the excision of the entire protein-coding sequence and a significant portion of the 5’ upstream region of the gene, respectively. Qualitative end-point PCR encompassing a 360 bp region between *ebony* exons e1 and e2 confirmed the absence of the gene in bp females from GUA10 (Fig. 2d). Altogether, these data show that the loss of the *ebony* coding sequence is responsible for the black pupae phenotype in *A*. *ludens* GUA10.

**Fig. 2:**
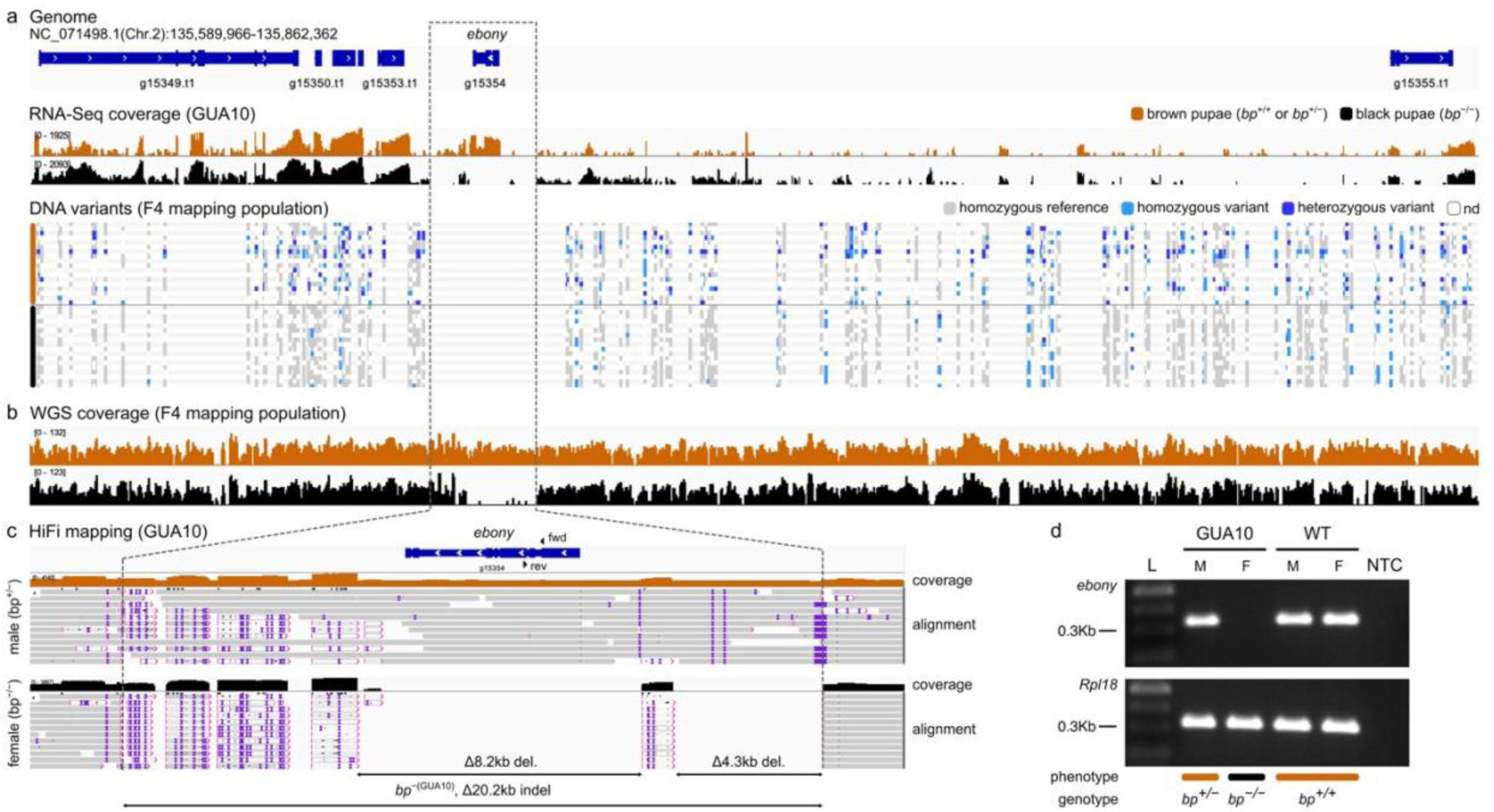
The bp phenotype in *A*. *ludens* GUA10 results from a large indel at the *ebony* locus. **a** Biallelic DNA variants between brown and black pupae individuals (*n* = 18 of each phenotype) from the F4 mapping population in the context of the *A*. *ludens* genome and GUA10 RNA-Seq read coverage (*n* = 3 males and females). No DNA variants are present within the *ebony* protein-coding sequence, coinciding with the absence of RNA-Seq reads from GUA10 females. **b** WGS read coverage (merged BAM files) from F4 population shows a large chromosomal interval with no coverage in black pupae individuals, suggesting the entire *ebony* coding sequence is missing in those flies. **c** Mapping of HiFi long-reads revealed a 20,182 bp DNA variant (NC_071498.1:g.135,665,293_135,685,474indel) within the *ebony* locus of *A*. *ludens* GUA10 (*bp*^−(GUA10)^), resulting in the removal of the entire protein-coding region of the gene. The variant is homozygous in black pupae female (*bp*^−/−^) and heterozygous in brown pupae male (*bp*^+/−^), as expected based on the genotypes of GUA10 GSS (Fig. 1a). **d** Qualitative end-point PCR of a 360 bp between *ebony* exons e1 and e2 confirmed the absence of this region in the GUA10 female genome (arrowheads in Fig. 2c indicate primer locations). Amplifications of *Rpl18* are internal controls. L = 100 bp DNA ladder (NEB), M = male, F = female, GUA10 = *A*. *ludens* GUA10 genetic sexing strain, WT = wildtype, and NTC = non-template negative control.

### Disruption of ebony creates analogous black pupae phenotypes in diverse tephritid flies

We used CRISPR/Cas9-mediated gene knockout (KO) to functionally validate our findings. Out of 260 *A*. *ludens* embryos injected with Cas9-sgRNA ribonucleoprotein (RNP) complexes targeting exon 1 of *ebony*, 38 (14.6%) survived into the pupal stage (Supplementary Table S3). Among them, 4 (10.5%) displayed the bp phenotype and developed into darker (melanic) adults (Fig. 3a), resembling the original mutant phenotype^13^ (*see* Fig. 1a). These mosaic knockout (mKO) flies contained indels in 56-89% of PCR-generated amplicons surround the sgRNA target site (Supplementary Fig. 3), confirming that the disruption of *ebony* is sufficient to induce the black pupae phenotype in *A*. *ludens*. Using a similar approach, we further recreated the mutant phenotype in *A. fraterculus* (Fig. 3b, Supplementary Fig. 4), demonstrating the functional conservation of *ebony* in other *Anastrepha* species.

**Fig. 3:**
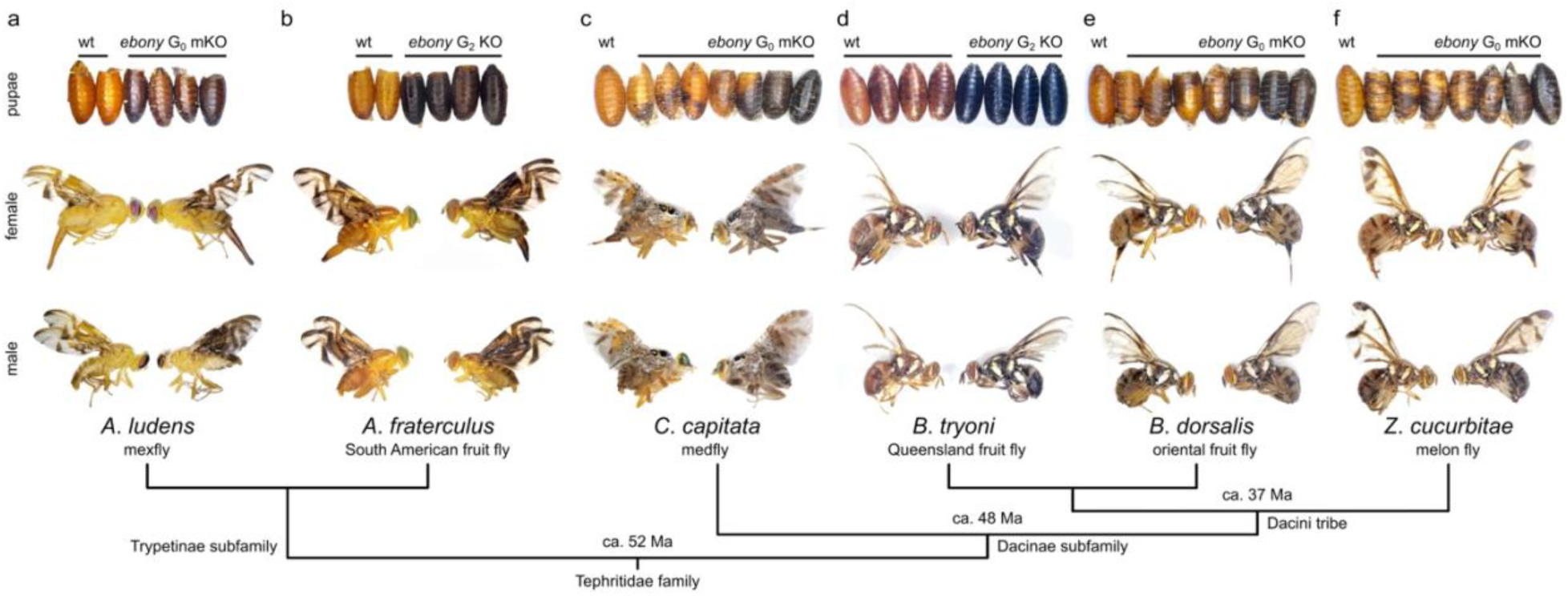
Disruption of *ebony* leads to the black pupae phenotype in diverse tephritids. **a** We used CRISPR/Cas9 to generate targeted loss-of-function mutations in the *ebony* gene, aiming to confirm its involvement in the bp phenotype of *A*. *ludens*. Surviving flies developed as either wildtype light-brown or mutant black puparia. Adults emerging from the latter also exhibited darker cuticles, mirroring the bp phenotype observed in GUA10. To assess the functional conservation of *ebony* in other tephritids, we extended our CRISPR experiments to include a close relative, *A*. *fraterculus* (**b**), as well as distantly related species, including *C*. *capitata* (**c**), *B*. *tryoni* (**d**), *B*. *dorsalis* (**e**), and *Z*. *cucurbitae* (**f**). Similar effects were observed in all species, demonstrating that the disruption of *ebony* is sufficient to induce the bp phenotype in diverse tephritids. wt = wildtype, mKO = mosaic knockout, KO = knockout, and Ma = million years. The evolutionary relationships among these species were estimated by Zhang et al.^24^ using mitochondrial phylogenomics.

Ebony is highly conserved among tephritids (Supplementary Fig. 5); thus, we anticipated that disrupting *ebony* orthologs would result in the bp phenotype even in distantly related species within the family (Fig. 3c-f). To test this assumption, we conducted additional CRISPR/Cas9 KO experiments on *C. capitata*, *B. dorsalis*, and *Z. cucurbitae*. Across all species, we observed high survival and mutagenesis rates (15-34% and 58-68%, respectively; Supplementary Table 3), and all experiments resulted in G_0_ individuals exhibiting phenotypes that ranged from mosaic black-brown puparium to complete melanized pupal cases (Fig. 3c, e, f). As expected by the nature of *bp* mutations in these flies^25–27^, the eclosed adults also displayed an overall darker cuticle. Genotyping of PCR products spanning Cas9 cut sites confirmed the presence of indels at the *ebony* loci, with modification frequencies ranging from 87-92% (Supplementary Fig. 6). Taken together, these results offered functional evidence supporting *ebony* as the responsible gene for the bp phenotype in mexfly GUA10 and likely other tephritids displaying parallel phenotypes.

### Ebony regulates pigmentation patterning in tephritid flies

Although mutations resulting in the bp phenotype are well-documented in tephritids, their impact on adult morphology has been overlooked. To address this, we established biallelic KO lines to investigate the contribution of *ebony* to adult pigment patterns. We focused on the conspicuous pictured wings of *C*. *capitata* and *Z*. *cucurbitae*, and the body pigmentation of *B*. *dorsalis* and *Z*. *cucurbitae*. These traits are prevalent among tephritids and thus may serve as ideal models for studying the evolution of pigmentation in this group of flies.

The wings of *C*. *capitata* exhibit intricate patterns featuring pigmented spots and bands varying in color and shape. Among these, the dominant discal crossband and the costal band are particularly intriguing (Fig. 4a), as their pigmentation is absent in model wings of *Drosophila* (examples in True et al.^28^). In *ebony* mutants, the appearance of those spots changed from brownish-yellow to dark brown; but their pattern remained the same (Fig. 4b). Interestingly, knocking out *ebony* had no effect on the silver lamina of medfly wings, while mutants of *Zeugodacus* displayed ectopic pigmentation in the structure (Fig. 4c, d). This ectopic pigmentation closely resembles that observed in *D. melanogaster*^23^, with no apparent changes in the apical spot, but a subtle enlargement of the fuscous black dm-cu crossvein blotch. A combination of modifications is observed in the wings of *A*. *fraterculus ebony* mutants (Supplementary Fig. 4). Similar to the medfly, the appearance of brownish-yellow bands changes to dark brown, and as in the melon fly, fuscous black edges intensify.

**Fig. 4:**
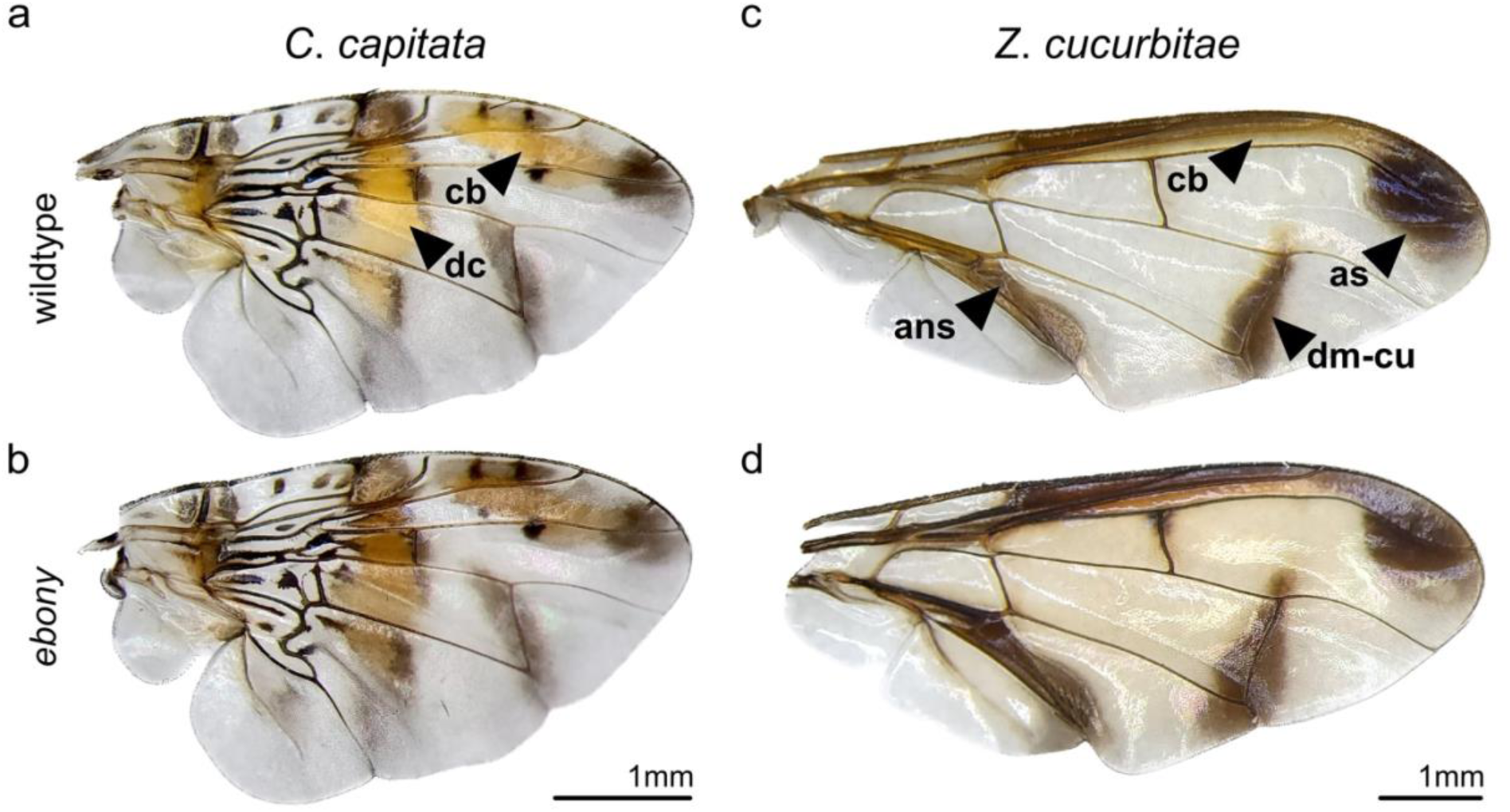
Ebony is required for proper wing pigmentation patterning in tephritids. **a** The wings of *C*. *capitata* are one of the most captivating examples of pictured pigmentation in tephritids. Among its intricate patterns, the dominant discal crossband (dc) and the costal band (cb) are particularly intriguing due to their uncommon yellowish-brown color. **b** In *ebony* mutants, the appearance of these bands changes to dark-brown, yet their pattern remains the same, with no observable effects on the lamina. **c** In contrast, the wings of *Z*. *cucurbitae* appear nearly clear, except for a few distinct pigmentation patterns, such as the dark apical spot (as) and the dm-cu crossvein blotch. **d** While a subtle enlargement of the dm-cu blotch appears in *ebony* mutants of *Z*. *cucurbitae*, the most striking phenotype is the ectopic pigmentation of the wing lamina, resulting in a smoky brown appearance. On an interesting note, the appearance of the anal streak (ans) and costal band (cb) in *Zeugodacus* wings also seems to change from dark-brown to black. Terminology in accordance to White and Elson-Harris^7^.

The *B*. *dorsalis* Punador strain maintained at the USDA-ARS-PBARC facility in Hawaii, exhibits a scutum predominantly black with peripheral reddish-brown areas (Fig. 5a). This pattern closely resembles the predominant morphotype found in the coastal areas of China^29^, which is the most likely source population that invaded the Hawaiian Islands^30^. In contrast, its abdomen is predominantly reddish-brown, displaying a distinct black T-shaped marking and narrow fuscous corners on tergites T4 and T5 (Fig. 5b, c). Loss of Ebony altered the pigment pattern in both body parts, allowing the breach of black melanin into areas where it was previously absent or constraint (Fig. 5d-f). These effects were even more dramatic in *ebony* mutants of the melon fly. The overall appearance of the thorax changed from orange-brown (Fig. 5g) to black (Fig. 5j), and the narrow abdominal bands (Fig. 5h, i) were significantly enlarged (Fig. 5k, l). These results illustrate the important role of Ebony in controlling the expressivity of black pigments in tephritids by inhibiting melanization, thus ensuring proper pigmentation patterning.

**Fig. 5:**
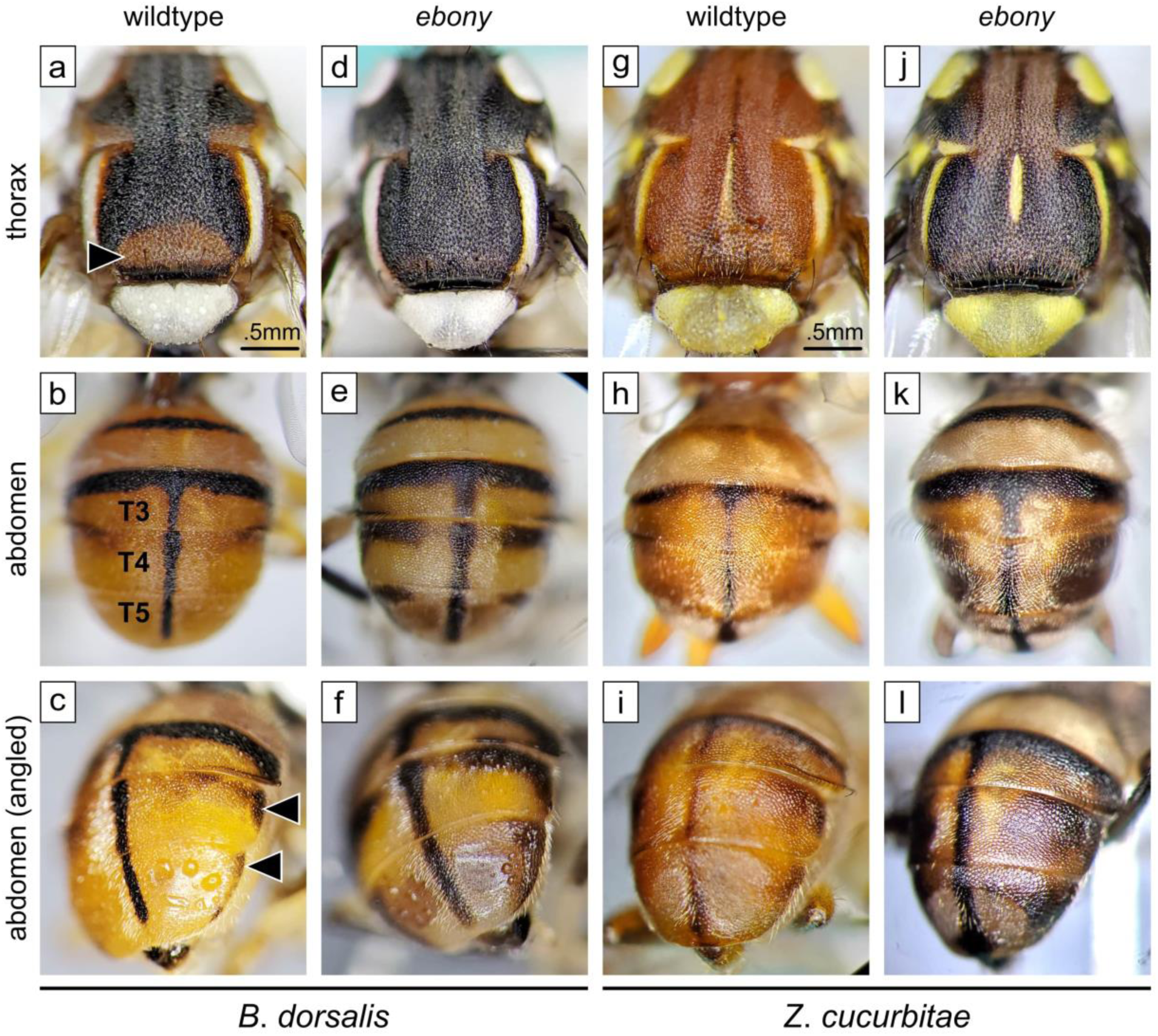
Ebony is required for proper body pigmentation patterning in tephritids. **a** Wildtype *B*. *dorsalis* (Punador strain) displays a predominantly black thorax with a particular reddish-brown area at the posterior edge of the scutum (arrowhead). **b**, **c** In contrast, its abdomen is predominantly reddish-brown with a distinct black T-shaped marking accompanied by narrow fuscous corners on tergites T4 and T5 (arrowheads). In *ebony* mutants, black pigmentation takes over the light areas of the thorax (**d**) while enlarging abdominal markings (**e**, **f**). Wildtype *Z*. *cucurbitae* displays a uniform light-brown color in the thorax (**g**) and abdomen (**h**, **i**), except for a few narrow black stripes in the latter. This overall appearance becomes completely distorted in *ebony* mutants. New pigmentation patterns appear as their thorax turns almost uniformly black (**j**), and their abdominal stripes become significantly enlarged (**k**, **l**). In both species, the absence of Ebony results in new pigmentation patterns. Terminology in accordance to White and Elson-Harris^7^.

### Development of an ebony-null strain for the Queensland fruit fly

The Queensland fruit fly *Bactrocera tryoni* is a significant agricultural pest in Australia^31^, where the SIT is applied to manage established populations and eliminate outbreaks in key horticultural regions. Presently, SIT targeting *B*. *tryoni* involves releasing both males and females, as no efficient method to remove females before sterilization exists. Therefore, creating a bp strain would represent a significant advancement in managing this important pest. To this end, we expanded our CRISPR/Cas9 experiments to KO *ebony* in *B*. *tryoni*. Out of 661 injected embryos, 12 survived to adulthood, all developing from brown pupal cases. We next backcrossed five G_0_ adults to the Ourimbah wt strain. All G_1_ flies were combined and allowed to interbreed. In total, 24 individuals with black puparium were recovered among the G_2_ offspring, confirming successful germline transmission of Cas9-induced mutations. Fourteen individuals survived to adulthood, all displaying dark cuticles (Fig. 3d). Molecular genotyping of G_2_ identified four *ebony* mutant alleles containing indels resulting in frameshift mutations (Supplementary Fig. 7). Initial attempts to develop a stable strain through inter-crossing G_2_ mutants failed due to non-viable eggs. However, we eventually established an *ebony*-null mutant strain (named *Bt*-*ebony*) after three rounds of backcrossing G_2_ homozygous mutants to the wt laboratory strain (*see* crossing scheme in Supplementary Fig. 8). Genotyping of *Bt-ebony* revealed a homozygous -2 bp deletion at *ebony* exon 1, resulting in a loss-of-function mutation as evidenced by the complete penetrance of the black pupae phenotype (Supplementary Fig. 7).

### Ebony has possible pleiotropic effects on the development of the Queensland fruit fly

We next asked if *ebony*-null alleles would result in fitness costs over the development of *B*. *tryoni*, an important consideration for potential SIT applications. We first conducted phenotype and genotype segregation analyses to test the effects on viability (Supplementary Table 4). *Bt-ebony* males (*e*^-/-^) were mass backcrossed to wildtype females (*e*^+/+^), resulting in hybrid offspring (*e*^+/-^) with brown pupae phenotypes. Inbreeding of F1 flies produced F2 progeny with wildtype (*n* = 200) or *ebony* (*n* = 60) phenotypes at an approximate 3:1 ratio (*X*^2^ = 0.51, *p* = 0.47, d.f = 1). Genotyping a subset of phenotypically wildtype F2 flies further confirmed the expected 1:2 segregation ratio of wildtype (*n* = 25) to heterozygous (*n* = 55) genotypes (*X*^2^ = 0.16, *p* = 0.69, d.f = 1), with no sex ratio bias. These results confirm the complete recessive inheritance of *ebony* and suggest that the generated mutant alleles do not impose viability costs on *B*. *tryoni*.

We further evaluated fecundity and development over three consecutive generations using four genetic crossing combinations: (1) wildtype control crosses; (2) test crossings with *Bt*-*ebony* males vs. wildtype females; (3) reciprocal crossings with wildtype males vs. *Bt*-*ebony* females; and (4) *Bt*-*ebony* crosses. All data were collected from F1 generations (Supplementary Table 5). For fecundity, we collected and counted eggs daily for five days. There were no significant differences in fecundity across all combinations. However, outcrossings (i.e., *Bt*-*ebony* vs. wildtype) and *ebony* crosses generally produced fewer eggs than wildtype control crosses (Fig. 6a). To assess hatchability, we counted eclosed larvae three days after egg collection. The percentage of hatched eggs from *ebony* crosses (median ± SE; 36.4 ± 5.2%) was significantly lower than in wildtype crosses (80.2 ± 5.5%; *p* = 0.04, one-way ANOVA Tukey HSD), while outcrossings produced intermediate values (Fig. 6b). Pupation, adult emergence, and partial adult emergence did not drastically differ between crossing combinations (Fig. 6c-e). Yet, *ebony* parents showed a tendency to produce offspring with more partially emerged adults (Fig. 6d, e). Additionally, the percentage of deformed adults was significantly higher among *ebony* progeny (3.86 ± 0.71%) compared to wildtype (0.58 ± 0.26%; *p* = 0.02, one-way ANOVA Tukey HSD; Fig. 6f).

**Fig. 6:**
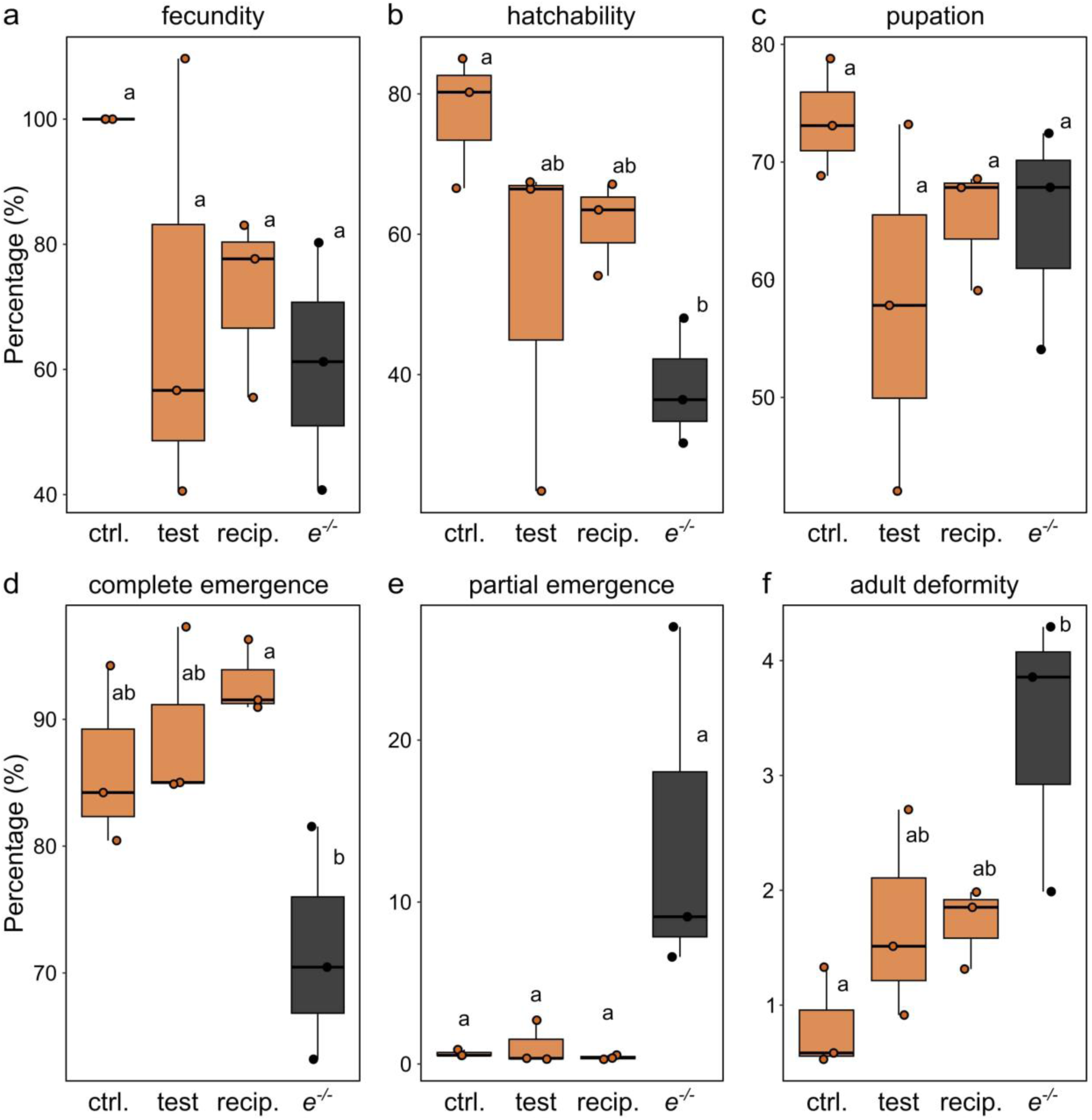
Ebony affects normal development in the Queensland fruit fly. We performed genetic crossings to evaluate whether ebony-null alleles would impose fitness costs over the development of *B*. *tryoni* expressed in terms of (**a**) relative fecundity (no. of eggs / no. of eggs in control experiments), (**b**) hatchability (no. of larvae / no. of eggs), (**c**) pupation (no. of pupae / no. of larvae), (**d**) adult emergence (no. of fully emerged adults / no. of pupae), (**e**) partial adult emergence (no. of partially emerged adults / no. of pupae), and (**f**) adult deformity (no. of deformed adults / no. pupae). Dots represent data collected from independent replicates (*n* = 3), box colors represent the pupae phenotype of their F1 offspring, and letters indicate significant differences (one-way ANOVA followed by Tukey’s HSD). ctrl. = control crossings between wt flies, test = test crossings between *Bt*-*ebony* males and wt females, recip. = reciprocal crossings between wt males and *Bt*-*ebony* females, and *e*^-/-^ = crossings between *Bt*-*ebony* flies.

## Discussion

In this study, we combined classical and modern genetics to uncover the molecular basis of the black pupae phenotype (bp) in tephritid fruit flies. We adopted the Mexican fruit fly *Anastrepha ludens* as our model—since stable bp mutants for the species exist as part of the GUA10 genetic sexing strain^14^. Genetic mapping and genome differentiation analysis allowed for the relative localization of the *black pupae* mutation (*bp*) within a ∼8.7 Mb interval on the mitotic chromosome 2 of *A*. *ludens*. Gene set enrichment analysis narrowed this causal region to two candidate loci: *ebony* and *yellow-f2*, which are key genes in the insect pigmentation pathway^21,23^. Comparative transcriptomics revealed that *ebony* expression is silenced in GUA10 females (homozygous recessive for *bp*) but not males (heterozygous for *bp*). This observation suggested that loss of *ebony* expression leads to the bp phenotype in GUA10 females, while the linkage of a functional allele to the Y-chromosome rescues the wildtype phenotype in males—since the bp is a recessive trait^13^. Supporting this model, cytogenetics of chromosomal translocations in GUA10 determined the locus responsible for the bp trait on the polytene chromosome 3 (mitotic chromosome 2) of *A*. *ludens*, and the fragment translocated into the Y-chromosome containing region 24^13,14^; the same region where we located *ebony* by *in situ* hybridization. DNA variant calling identified several small SNPs and indels surrounding *ebony* but none within its protein-coding region. A close examination of the short-read mapping revealed a large interval within the *ebony* locus lacking coverage from black pupae individuals of the mapping population, indicating the absence of the entire *ebony* gene in these flies. HiFi sequencing confirmed a 20,182 bp indel variant within the *ebony* locus, resulting in the removal of the entire protein-coding region of the gene. Finally, functional validation of *ebony* orthologues via CRISPR/Cas9-mediated KO generates analogous bp phenotypes in *A*. *ludens* and diverse tephritid species spanning over 50 Ma of divergent evolution. Collectively, these results provide strong evidence that *ebony* is the responsible gene for the bp phenotype in GUA10 and possibly other tephritids in which parallel phenotypes have been described. In addition to these findings, the creation of *ebony*-null mutants across distantly related species allowed for fundamental insights into the contribution of Ebony to color patterning and development in tephritids, which have significant importance for dipteran evolutionary developmental biology^19^, and thus discussed below. We hope that these thoughts will spark renewed interest in the evolution of pigmentation patterns and phenotypic diversity within the family Tephritidae.

### A pathway to black puparium

Ebony is crucial for insect pigmentation and sclerotization (hardening). It conjugates dopamine (DA) with β-alanine to produce N-β-alanyldopamine (NBAD), the precursor of yellow-tan sclerotin in insect cuticles^32–34^. Null mutants for *ebony* cannot produce NBAD, leading to an excess of DA, which in turn is diverted into the melanization branch of the pigmentation biosynthesis pathway (as reviewed by Massey and Wittkopp^35^), resulting in individuals with darker cuticles^32,36,37^. Consequently, a failure to encode Ebony leads to an increase in melanin (dark pigments) and a decrease in light pigmentation production^23^. In the classical example of *D. melanogaster*, *ebony* loss-of-function causes darkening of larval mouthparts and posterior spiracles, and adults much darker in appearance^22^. Our KO experiments yielded similar phenotypes, illustrating the functional conservation of Ebony across higher Diptera. Nonetheless, it’s worth mentioning a phenotypic difference between *ebony* mutants in *D*. *melanogaster* and the tephritids we studied. In the former, the loss of Ebony results in an unpigmented or pale pupal case^37^, while in the latter, it leads to an unusual black puparium. Naturally, the black pupae trait is not unique to tephritids, and similar phenotypes exist in *ebony* mutants of moths^38^, mosquitoes^39,40^, and—to some extent—other *Drosophila* species^41^.

The molecular mechanism driving this phenotypic divergence remains unknown (but *see* Sherald^37^), but it may involve the preferential utilization of DA during pupal cuticle formation, akin to the concept of strengths of the pathway (discussed by Spana et al.^42^). In insects, it’s accepted that N-acetyldopamine (NADA) serves as the primary precursor for colorless sclerotin, produced through the conjugation of DA by the *speck* gene. In contrast, brown cuticles predominantly originate from NBAD^33^. Thus, the NADA-to-NBAD ratio is expected to be a determinant factor for the final cuticle color intensity. Using the medfly as an example, NBAD is the primary precursor of sclerotization, giving rise to an opaque reddish-brown pupal case. In the absence of NBAD, as seen in *ebony* mutants, NADA is utilized instead^36,43^. However, the branch of the pigmentation pathway leading to NADA-sclerotin may not be as dominant as the one leading to NBAD-sclerotin. This difference in strengths would allow the conversion of excess DA into melanin, leading to the production of black pupal cases in *C*. *capitata*. In *D*. *melanogaster*, the branch leading to NADA-sclerotin is likely dominant, as evidenced by their clear, light-brown pupal cases. Therefore, this branch could be strong enough to prevent free DA from entering the melanization route in *ebony* mutants. Supporting this idea, the differential expression of *speck* may be solely responsible for the contrast between the black pupae of *Drosophila virilis* and the brown pupae of *Drosophila americana*^44^. Interestingly, it has been suggested that while DA can be directly converted into black melanin (mediated by Yellow), the conversion of DA into NBAD and then back into DA (mediated by Tan) seems necessary for the production of dark brown melanin^23,35^. The requirement for this indirect route could potentially explain why tephritid *ebony* mutants develop black pupae rather than darker brown ones. Finally, it should be noted that similar melanization phenotypes in *C*. *capitata* can also arise from the inability to produce β-alanine^43^, a condition that can be induced by mutations in the *black* locus^45^.

### Contributions of Ebony to wing pigmentation

Most tephritid flies are adorned with brightly contrasting body colors, and their wings often display elaborate patterns once described as both “lovely and mysterious”^46^. Wing pigmentation in tephritids can take various forms: from almost completely clear (*Bactrocera*), across relatively simple spots and blotches (*Zeugodacus*), and winding stripes shaded in black and brown (*Anastrepha*) to complex colored pictured wings with many patterns (*Ceratitis*). After a brief inspection, one could argue that such diversity is perhaps as enigmatic as in *Drosophila* and yet much less studied.

Here, we showed that Ebony is essential to generate and maintain this diversity. In *Zeugodacus*, Ebony inhibits melanization of the wing lamina, but it does not directly contribute to the pigmentation of the apical spot or crossvein blotches. Therefore, it is possible that Ebony is uniformly distributed in the species’ wings but less expressed in melanized patterns, and other genes (probably including *yellow*) are responsible for promoting their black pigmentation. This regulatory model is found in drosophilids exhibiting wing spots, where spatial downregulation of Ebony and upregulation of Yellow are—at least in part—responsible for their melanized patterns^23,47,48^. By masking melanization, Ebony also contributes to the expressivity of specific phenotypes, including the black crossvein blotch in *Z*. *cucurbitae* and the fuscous black edges of the S- and V-bands in the *Anastrepha* wings. Interestingly, ectopic pigmentation of the wing lamina, resembling that in *D*. *melanogaster ebony* mutants^23^, is also observed in *ebony* mutants of *Zeugodacus* and *Bactrocera* but not *Ceratitis* and *Anastrepha*. The mechanism behind this effect is unclear, but it seems to follow a phylogenetic pattern and therefore could be related to a species-specific distribution of pigment precursors through their wing veins.

The role of Ebony in the pigmentation of *C*. *capitata* wings was investigated by Pérez et al.^49^. Using biochemical assays, the authors showed that Ebony is spatially pre-patterned in the yellowish-brown bands of the medfly wings during early adult development, maintaining an extracellular activity long after epithelial cells disappear from the lamina. As observed in *Drosophila*^28^, pigment precursors present in hemolymph circulation gradually diffuse out from wing veins and are conjugated into NBAD by the pre-patterned Ebony to promote the coloration of these bands. Ebony activity is detectable in the yellowish-brown spots of *C*. *capitata* wing but not in the silver-paint lamina^49^, suggesting that the enzyme is only responsible for the pigmentation of the former. Our results provide functional evidence to their findings, showing that Ebony is necessary to produce the normal coloration of *C*. *capitata* wing bands and, in its absence, their appearance changes from yellowish-brown to dark brown—but their pattern remains the same, and no effects appear in the lamina. Notably, *Anastrepha ebony* mutants exhibit similar phenotypic changes in their wings, including the transformation of their once brownish-yellow S- and V-bands into a deep brown shade. The appearance of a dark-brown color in the wings of these *ebony* mutants can be explained by DA being shunted to the melanization branch of the pigmentation pathway and used to produce brown melanin through a mechanism open to debate. Some authors propose that the production of black and brown melanins occurs depending on the activity of different members of the Yellow gene family^50^. Others suggest that DA can be converted into brown melanin through a process involving phenol oxidases (POs; responsible for the oxidation of pigment precursors in the cuticle) but not the *yellow* gene^23,35^, and thus the indirect route must be taken (as discussed before). Our results seem to confront the latter idea, suggesting that brown pigmentation can be achieved through different molecular mechanisms in pupae and adults. These observations illustrate that the regulation of wing coloration in tephritid flies can be as mysterious as their pattern, and thus deserves more attention.

### Contributions of Ebony to adult body pigmentation

Natural populations of *B*. *dorsalis* exhibit extreme variation in adult pigment patterns^29,51^. This intraspecific diversity is particularly noticeable in their thorax (scutum), which ranges from pale reddish-brown to black, often with lanceolate-patterned intermediates. Similarly, their abdomen can vary from predominantly pale to predominantly black due to the expressivity of the typical ‘T’ pattern and the dark markings on tergites 4 and 5. The difference between morphological variants (morphs) is so dramatic that invasive populations of *B*. *dorsalis* in Africa were classified as a new species (*Bactrocera invadens*) and later synonymized^52^. Although the ecological relevance—if any—of these melanin patterns remains unknown, the geographic distribution of morphs seems to form irregular pigmentation clines^29^. The ectopic melanization in *ebony* mutants fairly overlap with some of these natural patterns, suggesting that this diversity in *B*. *dorsalis* may, in part, result from variations in the spatial expression of Ebony. The genetic background for ectopic expression also seems to contribute to the depth of these new patterns. For example, *ebony* mutants of the *B*. *dorsalis* Punador exhibit enlarged but distinct abdominal markings, which are nearly unrecognizable in predominantly black natural morphs^51^. In contrast, both thorax and abdominal segments of *ebony* mutants of the *B*. *tryoni* Ourimbah turn entirely black.

Within its genus, *Z*. *cucurbitae* is one-of-a-kind regarding adult morphology. Adults display a pale reddish-brown scutum and narrow abdominal bands, showing little intraspecific variability in natural populations^53^. These traits become completely distorted in *ebony* mutants; the scutum turns almost entirely black, and abdominal markings extend to a point where it resembles other related species (e.g., *Zeugodacus tau*). These shifts further illustrate how changes in Ebony expression may also confer phenotypic divergence between fruit fly species.

How do these phenotypic variations arise and how are they maintained? Based on our observations, and other more comprehensive studies^23^, we speculate these phenotypes are largely determined by the coordinated expression of Ebony and other pigmentation genes (such as *yellow* and *tan*), and changes in their expression patterns result in pigmentation diversity. Expression differences within and between species often arise from changes in either *cis*- or *trans*-acting elements important to these genes. In *Drosophila*, changes in *cis*-regulatory regions of *ebony* are responsible for the naturally occurring variation in abdominal coloration^54,55^ and, at some degree, to the intensity of the thoracic ‘trident’ phenotype^35^. Similarly, differences in expression levels of *ebony* and *tan* contribute to pigmentation divergence between the dark-bodied *Drosophila americana* and its light-colored sister species *Drosophila novamexicana*, which seems to be controlled in part by noncoding changes in these loci^56^. While divergence of pigment patterns within and among species might arise due to ecological and sexual selection pressures^3^, it remains uncertain whether it serves as raw material for speciation in species complexes with varying pigmentation, such as *B*. *dorsalis*.

### Potential pleiotropic effects of Ebony

The initial difficulties in establishing an *ebony* mutant line for *B*. *tryoni* could have been caused by off-target mutagenesis or genetic linkage. To mitigate and potentially eliminate these effects, we backcrossed Ebony mutants to wildtype flies for three consecutive generations before evaluating their relative fitness in terms of fecundity and development. Despite the effort, we found that *Bt*-*ebony* mutants exhibited a significant reduction in egg-hatching rates across three generations. Presumably, one disruption causes the other, suggesting that loss-of-function mutations in *ebony* impact traits other than pigmentation in the species. In insects, pleiotropy (a single gene affecting multiple traits) is often associated with pigmentation genes, and effects on behavior, physiology, and life-history traits have been extensively documented (reviewed by True^1^, Wittkopp and Beldade^2^, and Takahashi^57^). Although pleiotropic effects are difficult to prove, our results indicate that disruption of Ebony might compromise embryo viability in *B*. *tryoni*. Lower egg-hatching rates have also been reported in *ebony* mutants of the diamondback moth^58^ and silkworm^59^. In the latter, larvae seem to develop normally—but have trouble breaking out from their eggshell. It’s unclear whether *ebony* mutants of *B*. *tryoni* have impairments regarding larvae development or eclosion capabilities. Nevertheless, similar effects were observed in early adulthood, where *Bt*-*ebony* mutants appear to have difficulty emerging from their pupa, and a significant number of flies emerge with wing deformities. Interestingly, heterozygous individuals displayed intermediate egg-hatching and deformity rates, indicating semi-dominant phenotypes contrasting the apparent complete dominance of Ebony in pigmentation.

We also cannot discard the possibility of a higher frequency of copulation failure by *ebony* mutants, thus leading to an increased number of unfertilized eggs. Mutations in *ebony* have been correlated to abnormal mating behavior in *Drosophila*, likely due to defects in circadian rhythm and visual system (two well-known phenotypes of *ebony* mutants) and perhaps courtship behavior (for which ambiguous phenotypes exist)^57^. Furthermore, their relative abundance of cuticular hydrocarbons—that can function as conspecific short-range pheromones^60^—is also influenced by Ebony^41,61^. Additional studies, including more replicates and monitoring of mating trials, are needed to elucidate the influence of Ebony on the mating behavior of *B*. *tryoni*.

Further experiments are also needed to investigate whether additional backcrossings to wildtype could alleviate these deleterious effects and if the linkage of *ebony* to the male Y-chromosome could rescue them. In the same context, it’s reasonable to think that genome modifications at regulatory sequences could lead to the development of lines with fewer fitness costs—as regulatory mutations might minimize pleiotropic effects relative to coding mutations^1^. Therefore, there is a need to investigate the regulatory networks governing the spatiotemporal expression of pigmentation genes in tephritids. New technologies for profiling the transcriptome and epigenome at single-cell resolution (single-cell ATAC-Seq + RNA-Seq) could facilitate the identification of regulatory factors associated with their spatiotemporal expression. In theory, such investigation could help us manipulate the temporal expression patterns of these genes, thereby bypassing the downsides associated with their disruption. Fitness costs are also likely to arise from random radiation-induced mutagenesis. Potentially, this could be overcome by CRISPR/Cas9-mediated knock-in or chromosomal translocation of the rescue allele into the Y-chromosome—as previously proposed^17,62^—for which standardized protocols remain to be established.

## Methods

### Flies

*Anastrepha ludens* flies (wildtype) used for microinjections were obtained from the USDA-APHIS Moore Air Base Facility in Edinburg, TX, USA. Flies are reared at 23 °C and 60% RH under a 14/10 h light/dark cycle. Adults are fed with a standard dry diet (1 yeast: 3 sugar) and water. Larvae are reared on a meridic diet consisting of 15.6% pelletized corncob, 8.4% granulated white sugar, 2.8% torula yeast, 7% wheat germ, 4.6% toasted soy flour, 2.8% corn flour, 0.2% vitamin mix, 1.9% citric acid, 0.1% sodium benzoate, 0.2% methylparaben, and 0.2% bravo WS (1%), all dissolved in water. Samples of the *A*. *ludens* GUA10 strain and F4 mapping populations, used for WGS, RNA-Seq, and HiFi sequencing, were obtained from the USDA-APHIS MOSCAMED San Miguel Petapa Fruit Fly Rearing and Quarantine facility in San Miguel Petapa, Guatemala.

*Anastrepha fraterculus* sp. 1 flies (Vacaria strain) were obtained from the Insect Pest Control Laboratory in Seibersdorf, Austria. Flies are maintained at 25 °C and 48% RH under a 12/12 h light/dark cycle. Adults are fed with a standard dry diet (1 yeast: 3 sugar) and water. Larvae are reared on a carrot diet consisting of 8.06% brewer’s yeast, 0.76% sodium benzoate, 0.8% (v/w) HCl, 23.9% carrot powder, and 67.17% fresh carrots, all dissolved in water.

*Ceratitis capitata* (rearing strain), *Bactrocera dorsalis* (Punador strain), and *Zeugodacus cucurbitae* (rearing strain) were obtained from the USDA-ARS-PBARC in Hilo, HI, USA. All species are maintained at 25 °C and 60% RH under a 12/12 h light/dark cycle. Adults are fed with a standard dry diet (1 yeast: 3 sugar) and water. Larvae are reared on a species-specific diet. The *C*. *capitata* larval diet consists of 59% wheat mill feed, 27.3% granulated white sugar, 7.6% torula yeast, 5.2% citric acid, and 0.5% each of the preservatives nipagen and sodium benzoate. The *B*. *dorsalis* larval diet consists of 63% wheat mill feed, 28.5% granulated white sugar, 8% torula yeast, 0.3% nipagen, and 0.2% sodium benzoate. The *Z*. *cucurbitae* larval diet consists of 73.7% wheat mill feed, 17.4% granulated white sugar, 8.4% torula yeast, and 0.3% each of the preservatives nipagen and sodium benzoate. All diets are dried and mixed with water before being provided to larvae.

*Bactrocera tryoni* flies (Ourimbah strain) were obtained from the New South Wales Department of Primary Industries in Ourimbah, Australia. Flies are reared at 25 °C and 65% RH under a 14/10 h light/dark cycle. Adults are fed with a standard dry diet (1 yeast: 3 sugar) and water. Larvae are reared on a gel diet containing 20.4% brewer’s yeast, 12.1% sugar, 0.2% methyl p-hydroxy benzoate, 2.3% citric acid, 0.2% wheat germ oil, 0.2% sodium benzoate, and 1% agar.

### Mapping population

The crossing scheme shown in Fig. 1b was used to introgress the mexfly *black pupae* mutation into a wildtype reference genetic background and generate a mapping population for genome-wide genetic differentiation analysis. Briefly, mexfly black pupae females were isolated from the GUA10 strain^14^ during pupal stage and individually outcrossed to wildtype males to produce hybrid offspring with non-sex-linked variation in pupal color. The resulting F1 population displayed the wildtype brown pupae phenotype and was allowed to intercross freely to recover the black pupae phenotype in the next generation. Resulting F2 black pupae females were individually backcrossed to wildtype males. Heterozygous flies at F3 were let to inbreed, and black and brown pupae siblings were selected at F4. Individuals were separated by pupae color, snap frozen in liquid nitrogen upon adult emergence, fixed in absolute ethanol and kept at -80 °C until further analysis.

### Association mapping

Total DNA was extracted from brown (*n* = 18) and black (*n* = 18) pupae individuals of the F4 mapping population using the NucleoMag Tissue kit (Macherey Nagel) on a Kingfisher Flex system. Extractions were quantified by Qubit dsDNA BR assay (Invitrogen), normalized to 50 ng on a 96-well plate, and subjected to library preparation using the RipTide High Throughput Rapid DNA Library Preparation kit (iGenomX). Libraries were pooled and sequenced on a single lane of HiSeq 4000 (Illumina) in a 150 bp paired-end run. Raw data was demultiplexed and processed in-line to remove barcodes using the fgbio toolkit (https://github.com/fulcrumgenomics/fgbio).

Demultiplexed whole-genome sequencing (WGS) reads were then filtered using fastp v0.23.2^63^ and mapped against the mexfly reference genome using BWA-MEM v.2.2.1^64^. SAMtools v1.17^65^ *fixmate* was used to update mate information and duplicated pairs were marked using SAMBLASTER v0.1.26^66^. Alignments were filtered using SAMtools *view* with the following parameters: skip alignments with mapping quality ≤ 30, only keep reads mapped in a proper pair (-f 0x0002), and discard unmapped reads (-F 0x0004), reads with unmapped mates (-F 0x0008), and reads marked as duplicates (-F 0x0400). Genome Analysis Toolkit v4.4 (GATK, https://gatk.broadinstitute.org/) *RealignerTargetCreator* and *IndelRealigner* were used to locally realign reads around insertions and deletions (InDels) in filtered alignments. Variant calling was performed using GATK *HaplotypeCaller*, *CombineGVCFs*, and *GenotypeGVCFs* with default parameters. Variants were hard filtered using GATK *SelectVariants* and *VariantFiltration* following the recommended parameters for SNPs and InDels to pass. Remaining variants were further filtered using VCFtools v0.1.16-9^67^ with the following parameters: maf 0.05, mac 2, min-alleles and max-alleles 2, min-meanDP 0.2 and max-meanDP 1.8 (based on 5th and 95th percentiles of mean depth per site), hwe 1e-5, max-missing 0.5, and minQ 30. Genome-wide genetic differentiation index (F_ST_) between black and brown pupae siblings from the mapping population were calculated in 100 Kb windows at 20 Kb sliding intervals using VCFtools and visualized with the *qqman* package in R v4.3.2. The final, filtered VCF (Variant Call Format) file was examined in the Integrative Genomics Viewer (IGV) tool v2.16.2-0^68^.

### Genome annotation

RepeatModeler v2.0.4^69^ was used to identify repetitive sequences and construct a *de novo* repeat library for the *A*. *ludens* reference genome. This library was combined with *Drosophila* repetitive sequences extracted from RepBase (RepeatMasker Edition v.20181026, https://www.girinst.org/server/RepBase/index.php) and used with RepeatMasker v4.1.5 (https://www.repeatmasker.org/) to mask repetitive sequences in the genome. Gene prediction was carried out using RNA-Seq alignment data as extrinsic evidence for gene models. Publicly available RNA-Seq raw reads, encompassing most of *A*. *ludens* development stages (Supplementary Table 6), were filtered using fastp v0.23.2^63^ and mapped to the masked genome with STAR v2.7.10b^70^ in two-pass spliced alignment mode. The resulting alignments were used as input for BRAKER2 pipeline^71^. The longest isoform of each predicted gene model was retrieved using AGAT v0.8.0 (https://github.com/NBISweden/AGAT) function *agat_sp_keep_longest_isoform.pl* resulting in an initial set of 18,230 protein coding genes. BUSCO v5.4.5 analysis^72^ found 96.9% of the 3,285 single-copy orthologues (in the diptera_odb10 database) to be complete (96.3% single-copied and 0.6% duplicated genes), 0.8% fragmented, and 2.3% missing. Functional annotations were performed by mapping this gene set against the UniProtKB database (https://www.uniprot.org/) using BLASTp (*e*-value ≤ 1e-6), and the Pfam, PANTHER, Gene3D, SUPERFAMILY, SMART, and CDD databases using InterProScan v5.64-96.0^73^. Orthology assignments were performed with the DIAMOND (*e*-value ≤ 1e-5) mapping mode implemented in eggNOG-mapper v2.1.12-0^74^ in the context of Diptera taxonomic scope. The final gene set for *A*. *ludens* genome contained 13,114 full-length protein-coding genes with annotated gene ontology (GO) terms.

### Enrichment analysis

Gene ontology (GO) enrichment analysis of biological processes terms was performed using the Bioconductor package topGO v2.54.0-0 (https://bioconductor.org/packages/topGO) on the gene set within the black pupae causal region on mexfly chromosome 2, using all genes with annotated GO terms in the genome as the background set. The *elim* method was used to reduce redundancy and searches were limited to categories containing at least 10 annotated genes. Overrepresentation significance of GO terms was calculated using the classical Fisher test adopting a cutoff threshold of *p*-value ≤ 0.05.

### RNA isolation

Specimens were euthanized at -20 °C, fixed in RNAlater (Invitrogen) and stored at 4 °C until further processing. Before extractions, samples were mixed with one volume of 1x PBS (pH 7.4) and centrifuged at top speed for 10 min at 4 °C, allowing removal of solution excess by pipetting. Pre-processed samples were homogenized in TRISure (Bioline Meridian Bioscience), and total RNA was isolated using Direct-zol-96 MagBead RNA kit (Zymo Research). Extractions were DNase treated in a 50 µL reaction containing 2 U of TURBO DNase (Invitrogen), 50 U of RiboGuard RNase Inhibitor (Lucigen), 1x TURBO reaction buffer and 5-10 µg of total RNA. Reactions were performed at 37 °C for 30 min, stopped by the addition of 15 mM of EDTA (pH 8.0) followed by an incubation at 75 °C for 10 min, and purification using the RNA Clean & Concentrator-5 kit (Zymo Research).

### RNA-Seq

RNA sequencing (RNA-Seq) libraries were generated from 250 ng of DNase-treated RNA with the NEBNext Ultra II Directional RNA Library Prep Kit for Illumina (NEB) following the Poly(A) mRNA Magnetic Isolation Module protocol. Sequencing was performed on a AVITI platform (Element Biosciences) in a 150 bp paired-end run. Raw reads were filtered using fastp v0.23.2^63^ and mapped against the *A*. *ludens* reference genome with STAR v2.7.10b^70^ using the two-pass spliced alignment mode along with gene models generated by BRAKER2. Mapped reads were quantified by featureCounts^75^ implemented in subread v2.0.4 (https://subread.sourceforge.net/) and resulting count matrix used as input for differential expression analysis with the Bioconductor package edgeR v4.0.12^76^. Briefly, counts were first filtered by the *filterByExpr* function and normalized using the TMM-method (Trimmed mean of *M*-values). Differences between brown (*n* = 3) and black (*n* = 3) pupae samples were estimated with the quasi-likelihood (QL) F-test against the threshold of log_2_ fold-change (logFC) > 2 using the *glmTreat* function. Genes were considerate differentially expressed when false discovery rate (FDR) < 0.05. Read counts were converted to RPKM (reads per kilobase of transcript per million reads mapped) with the *rpkm* function and used as a descriptive measure of gene expression for visualization.

### HiFi long-read mapping

High molecular weight (HMW) DNA was extracted from single adult flies using the MagAttract HMW DNA Kit (QIAGEN). Samples were quantified using the Qubit dsDNA BR assay (Invitrogen) and size checked in the Femto Pulse System (Agilent Technologies). Extractions were further subjected to a 2x bead cleanup, and their purity determined on the basis of OD 260/230 and 260/280 ratios estimated in a DS-11 spectrophotometer (DeNovix). Clean HMW DNA samples were sheared to a mean size of 20 kb using the Megaruptor 2 (Diagenode), and size checked on the Fragment analyzer (Agilent Technologies) with the HS Large Fragment kit. SMRTBell libraries were prepared using approximately 1 µg of sheared DNA with the SMRTBell Prep Kit 3.0 (Pacific Biosciences). The prepared libraries were bound and sequenced on a 24M SMRT Cell on a Revio system (Pacific Biosciences) using a 24 h movie collection time. Circular consensus sequences were obtained using SMRTLink v13.0 on the Revio instrument. Highly accurate long-reads (HiFi reads) were filtered for adapter contamination with HiFiAdapterFilt v.3.0.1^77^ and mapped to the *A*. *ludens* reference genome with minimap2 v.2.24^78^ (-ax map-hifi). Mapping files were examined in the IGV tool v2.16.2-0^68^.

### Manual gene annotation

Gene models for *ebony* orthologues (Supplementary Data 1) were manually curated by mapping the *D*. *melanogaster* Ebony peptide sequence (FlyBase: FBgn0000527) against each reference genome using tBLASTn (BLOSUM45 matrix, *e*-value ≤ 1e-5). Highly similar genomic regions were retrieved with BEDTools v2.31.1-0^79^ and intron-exon boundaries modelled using Exonerate v2.4.0-7^80^ protein2genome mode (--refine full). Intron-exon boundaries were manually annotated, coding sequences were translated using the ExPASy translate tool (https://web.expasy.org/translate/), and signature motifs identified using DELTA-BLAST (https://blast.ncbi.nlm.nih.gov/Blast.cgi) against the NCBI’s non-redundant protein database.

### Semiquantitative RT-PCR and qualitative end-point PCR

For RT-PCRs, first-strand cDNAs were synthesized from 1 µg of DNase-treated RNA using the MMLV Reverse Transcriptase Synthesis Kit (Lucigen) protocol with the Oligo(dT)_21_ primer. Reverse transcriptions were performed at 37 °C for 1 h, terminated at 85 °C for 5 min and stored at -20 °C. Reverse Transcription PCRs (RT-PCR) were carried out for a final volume of 12.5 µL containing 1x One*Taq* HS Quick-Load Master Mix with Standard Buffer (NEB), 0.2 µM of each forward and reverse Intron-flanking primers (Supplementary Table 7), and an equivalent amount of cDNA synthetized from 50 ng of total RNA. Amplifications were carried out in the following conditions: 94 °C for 3 min, 40 cycles of [94 °C for 30 s, 55 °C for 30 s and 68 °C for 1 min], and a final extension at 68 °C for 5 min. All experiments included independent biological (*n* = 3) and technical (*n* = 3) replicates, no template controls (cDNA omitted from the reaction and volume adjusted with nuclease-free water), and a genomic DNA control (total DNA extraction from a single *A*. *ludens* wildtype individual). Amplifications were resolved in 1% agarose gels stained with Midori Green Advance DNA Stain (at 1:15000 dilution factor; Nippon Genetics) in 0.5x TBE buffer. Images were acquired in a Gel Doc XR+ Gel Documentation System (Bio-Rad) using predefined setups in the software Image Lab v.6.0.1. Contrast and light corrections were made in the same software. For end-point PCRs, genomic DNA was extracted from whole adult flies using the NucleoMag Tissue Kit (Macherey-Nagel), and a total of 50 ng was used as a template for PCR amplifications encompassing *ebony* exons e1 and e2 with One*Taq* HS Quick-Load Master Mix with Standard Buffer (NEB) as described before.

### Cytogenetics

Third-instar larvae salivary glands of wildtype *A*. *ludens* were dissected in 45% acetic acid and fixed in glacial acetic acid: water: lactic acid (3:2:1) solution for about 5 min. Preparations were stored overnight at -20 °C and dipped into liquid nitrogen in the next day. Slides were dehydrated in absolute ethanol, air dried, and stored at 4 °C. DNA probes for fluorescent *in situ* hybridization (FISH) of *ebony* were prepared by PCR for a final volume of 25 µL containing 1x Platinum II Hot-Start Green PCR Master Mix (Invitrogen), 0.2 µM of each forward and reverse primer (Supplementary Table 7), and 100 ng of total DNA. Amplifications were carried out as follows: 94 °C for 5 min, 35 cycles of [94 °C for 45 s, 56 °C for 30 s and 72 °C for 1.5 min], and a final extension at 72 °C for 1 min. Probe labeling was performed according to the DIG DNA Labelling Kit (Roche) protocol, and slides were prepared for fluorescence detection as previously described^17^. Hybridizations in isolated chromosomes were photographed in a Leica DM2000 LED microscope using a Leica DMC5400 digital camera and analyzed with the LAS X software v3.7.0 in the context of the mexfly salivary gland chromosome maps^81^.

### CRISPR/Cas9

Knockout experiments in *A*. *ludens*, *C*. *capitata*, *B*. *dorsalis*, and *Z*. *cucurbitae* were performed according to a proposed standard protocol for Cas9-mediated gene disruption in non-model tephritids^62^. Briefly, single guide RNAs (sgRNAs) were designed against the *ebony* gene using CRISPOR v.5.01^82^ and their reference genomes (Supplementary Table 7). Templates for sgRNAs were generated by PCR with Phusion High-Fidelity DNA Polymerase (NEB), purified using the QIAquick PCR purification kit (QIAGEN), and used for T7 *in-vitro* transcription with MEGAshortscript Kit (Invitrogen). Purified Cas9 protein conjugated with a nuclear localization signal (NLS) was obtained commercially (PNA Bio). Microinjection mixes contained end-concentrations of 360 ng/µl Cas9, 200 ng/µl sgRNA, 1x injection buffer (0.1 mM Sodium phosphate buffer, 5 mM KCl), and 300 mM KCl in a final volume of 10 μL in nuclease-free water. Ribonucleoprotein (RNP) complexes were pre-assembled at 37 °C for 15 min and delivered through the chorion into the posterior end of preblastoderm embryos within the first 60 min after egg-laying (AEL). Mosaic black-brown puparia at G_0_ were inbred to establish biallelic mutant lines.

Microinjections in *A*. *fraterculus* followed the same protocol with minor modifications. The sgRNA was designed using CHOPCHOP v.3^83^ in combination with Geneious v.2023.0.2 (https://www.geneious.com/) in the context of *A*. *fraterculus* draft genome. Both sgRNA (Merck) and purified Cas9 protein (PNA Bio) were obtained commercially. RNPs were pre-assembled at 37 °C for 15 min, followed by 5 min at room temperature. Embryos were collected every 30 min, dechorionated with a 30% hypochlorite solution for 80 s, and microinjected within 120 min AEL. Injected G_0_ flies were individually backcrossed to wildtype flies. The resulting G_1_ offspring were inbred, and all G_2_ displaying the bp phenotype were intercrossed. Adult individuals at G_3_ were genotyped using non-lethal methods, and flies harboring the -7 bp deletion allele were inbred to establish a homozygous mutant strain at G_4_.

For knockout experiments in *B*. *tryoni*, purified Cas9 protein (IDT Alt-R S.p. Cas9 Nuclease 3NLS) and guide RNAs (IDT customized Alt-R crRNAs and universal Alt-R tracrRNA) were obtained commercially. Two customized crRNAs were designed using CRISPOR v.5.01^82^ in the context of the species reference genome and a multiple sequence alignment of target regions to avoid polymorphisms in the injection population. Two dual-guide RNA (dgRNA) duplexes were annealed separately by mixing 40 μM of each specific crRNA with 40 μM of universal tracrRNA in Nuclease-Free Duplex Buffer and heating at 95 °C for 5 min before cooling to room temperature. The final injection mix contained 300 ng/µL Cas9 protein, 10 μM of each dgRNA, and 1x injection buffer (0.1 mM sodium phosphate buffer pH 6.8, 5 mM KCI) in a final volume of 10 µL in Nuclease-Free Duplex Buffer. The injection mix was incubated at room temperature for 5 minutes to allow RNP formation and delivered into the posterior end of embryos within 60 min AEL. Surviving G_0_ adult flies were individually backcrossed to wildtype flies, and their G_1_ progeny were combined and allowed to interbreed. Biallelic *ebony* mutants were recovered at G_2_ and used to stablish the *Bt-ebony* homozygous mutant strain (Supplementary Fig. 7, 8).

### Genotyping

Mosaic knockouts (mKO) of *A*. *ludens*, *C*. *capitata*, *B*. *dorsalis*, and *Z*. *cucurbitae* were genotyped by deep-sequencing of indexed amplicons surrounding the Cas9 cut sites, as previously described^62^. Briefly, total DNA was extracted from whole adult flies using the NucleoMag Tissue Kit (Macherey-Nagel) and used as a template for a two-step PCR with Phusion High-Fidelity DNA polymerase (NEB) and primers in Supplementary Table 7. Indexed amplicons were purified with the QIAquick PCR purification kit (QIAGEN), pooled in equimolar ratios, and sequenced on Illumina iSeq 100 system (150 bp paired-end reads). Targeted genome modifications were inspected using CRISPResso2 v2.2.14-0^84^. Biallelic KO mutants examined in phenotypic analysis were genotyped in the same way.

Total DNA from whole G_2_ adults of *A*. *fraterculus* was extracted using the ExtractMe Genomic DNA kit (QIAGEN) and used as a template for PCR amplifications with Platinum Green Hot Start PCR Master Mix (Invitrogen) and primers in Supplementary Table 7. Reactions were cycled as follows: 94 °C for 2 min, 34 cycles of [94° C for 15 s, 59 °C for 15 s, and 68 °C for 15 s], and a final extension at 68 °C for 5 min. Amplification products were purified using the ZR-96 DNA Clean-up kit (Zymo Research) and Sanger sequenced on an ABI 3730XL DNA Analyzer system (Applied Biosystems). Site-specific modifications were inspected in a multiple sequence alignment produced by Geneious v.2023.0.2. Non-lethal genotyping of G_3_ flies was carried out using single adult legs with Platinum Direct PCR Universal Master Mix (Invitrogen).

Amplification of DNA surrounding the Cas9 cut sites in *B*. *tryoni* was performed directly from single adult legs using the Phire Animal Tissue Direct PCR Kit (Thermo Scientific) and primers in Supplementary Table 7. Amplification conditions were as follows: 98 °C for 5 min, 35 cycles of [98 °C for 5 s, 60 °C for 5 s, and 72 °C for 20 s], and a final extension at 72 °C for 1 min. For the identification of heterozygous G_2_ brown pupal flies, amplifications were submitted to cleavage assays using the T7 Endonuclease 1 (T7E1) enzyme protocol (NEB). PCR products of confirmed G_2_ heterozygous (*n* = 48) and final *ebony*-null mutants (*n* = 30) were Sanger sequenced on an ABI 3730XL DNA Analyzer system (Applied Biosystems). Targeted genome modifications were inspected in a multiple sequence alignment produced by Geneious v.2023.0.2.

### Performance assays

Segregation analysis was performed to measure the viability of *B*. *tryoni* individuals carrying *ebony*-null alleles. To account for environmental variations, data were collected after the 1^st^ and 2^nd^ rounds of mass backcrossing between *ebony* males and wildtype females during the establishment of the *Bt-ebony* strain (Supplementary Fig. 8). All F1 individuals were interbred, and phenotypic segregation at F2 was tested against the 3:1 Mendelian inheritance ratio of phenotypes. A subset of phenotypically wildtype F2 flies were further genotyped by T7E1 assays, and the segregation of wildtype to heterozygous was tested against a 1:2 inheritance ratio of genotypes. Pearson’s chi-square goodness-of-fit tests were used to determine significant deviation from expected ratios as implemented in the *stats* package in R v4.3.2.

Fitness analysis was performed to measure the relative fecundity and development of *B*. *tryoni ebony* mutants. Virgin females and naive males from the *Bt-ebony* and wildtype Ourimbah strains were sorted within three days after adult emergence and kept separated until experimentation. The following crosses were performed using 8-to-11-days-old flies (10 males and 10 females): (1) wildtype males vs. wildtype females; (2) *ebony* males vs. wildtype females; (3) wildtype males vs. *ebony* females; and (4) *ebony* males vs. *ebony* females. Assays were performed over three consecutive generations to account for environmental variations (Supplementary Fig. 8). The first batch of eggs was collected 48 h after experiment setups. Eggs were collected for 24 h, and the number of eggs laid was counted daily for 5 days to assess fecundity ratios. Collected eggs were placed on larval diet and resulting larvae were counted 3 days after incubation to estimate embryonic hatching rates. The number of larvae that reached the pupal stage was used to estimate pupariation rates. Emerging adults were recorded daily until no new eclosions were observed and categorized as fully emerged, partially emerged (remaining in the puparium), and deformed (emerged flies with twisted or deformed wings). Significant differences between groups at *p* ≤ 0.05 were determined by one-way ANOVA followed by a Tukey’s Honest Significant Difference (HSD) tests using the *stats* package in R v.4.3.2.

## Data availability

The following reference genome assemblies were used in this study: *A*. *ludens* (GenBank: GCA_028408465.1), *C*. *capitata* (GenBank: GCA_000347755.4), *B*. *dorsalis* (GenBank: GCA_023373825.1), *B*. *tryoni* (GenBank: GCA_016617805.2), and *Z*. *cucurbitae* (GenBank: GCA_028554725.2). All raw sequencing data generated in this study are deposited in the NCBI Sequence Read Archive (SRA) database under the BioProject PRJNA1139181. SRA accession numbers for each sample are detailed in Supplementary Table 6. Final de novo genome annotation files for *A*. *ludens* are archived on figshare repository at https://doi.org/10.6084/m9.figshare.26376841. Manually curated annotations of *ebony* orthologues can be found in Supplementary Data 1. All primers used in this study are listed in Supplementary Table 7. The source data for fitness analysis of the Bt-ebony strain is available as Supplementary Table 5.

## Acknowledgements

The authors would like to thank N. Barr, E. Braswell, T. Todd, H. Conway, K. Pina, and A. Arellano for their support with microinjections and rearing of *A*. *ludens* at the USDA-APHIS Moore Air Base Facility (Texas, USA). We thank D. Rubinoff and C. Doorenweerd for their insights into the adult color patterns of *B*. *dorsalis*. We would also like to acknowledge C. Sylva for her assistance in fly rearing at the USDA-ARS-PBARC (Hawaii, USA), M. Okamoto for confirming *Bt*-*ebony* genotypes, and A. Spengler for the scientific illustrations displayed in Fig. 1b. This research was supported by the in-house appropriated USDA-ARS project *Advancing Molecular Pest Management, Diagnostics, and Eradication of Fruit Flies and Invasive Species* (no. 2040-22430-028-000-D), the USDA Plant Protection Act Section 7721 project *Develop Techniques to Rapidly Generate Non-Transgenic Female Lethal Genetic Sexing Strains in Tephritid Pests* (no. 6.0179), the HSF 18-6 Hermon Slade Foundation grant, and the Insect Pest Control Subprogramme of the Joint FAO/IAEA Centre of Nuclear Techniques in Food and Agriculture. This study also benefitted from discussions at meetings for the Coordinated Research Project *Generic Approach for the Development of Genetic Sexing Strains for SIT Applications* (no. D44003), funded by the International Atomic Energy Agency (IAEA), and used resources provided by the SCINet project of the USDA-ARS projects no. 0201-88888-003-000D and 0201-88888-002-000D. USDA is an equal opportunity provider and employer. Mention of trade names does not imply an endorsement from USDA or the Federal Government.

## Contributions

The initial evidence linking *ebony* to the black pupae phenotype in the *A*. *ludens* GUA10 strain was independently conceived by DFP and SMG at the USDA-ARS-PBARC (Hawaii, USA) and by CMW, TNMN and SWB at the University of Melbourne (Melbourne, AUS). This work combines and expands the data of these two preliminary studies. DFP, TNMN, CMW, AC, PC, KB, SWB, SBS and SMG conceptualized this study. SBS and SMG designed and supervised the establishment of the *A*. *ludens* F4 mapping population in Guatemala. PR and REYR established the mexfly mapping population, collected and pre-processed samples for WGS and RNA-Seq. RLC prepared samples and libraries for WGS, RNA-Seq, and HiFi sequencing with the assistance of ANK. DFP performed the bioinformatics analyses with insights from SBS and SMG. GG performed the *in situ* hybridization of *ebony* in *A*. *ludens* chromosomes. DFP performed microinjections in *A*. *ludens* with support from ANK. CRISPR experiments in *A*. *fraterculus* were made by AASC, and strains were maintained by AASC. DFP performed CRISPR experiments in *C*. *capitata*, *B*. *dorsalis*, and *Z*. *cucurbitae*. DFP established and maintained KO strains with the assistance of ANK. TNMN performed CRISPR experiments in *B*. *tryoni*. The *Bt*-*ebony* mutant strain was established and maintained by AO, AC, EF and TNMN. TNMN, AO, AC, and EF performed segregation and fitness analysis of *Bt-ebony* mutants. PC, KB, SBS, SWB, and SMG supervised, administered, and/or secured funding for this work. The same authors assisted with data interpretation and manuscript revision. DFP wrote the manuscript with substantial insights from the initial draft written by TNMN. All authors read and approved the final version of the manuscript.

