## Supplementary for "The genetic basis of the black pupae phenotype in tephritid fruit flies"

by

Paulo and Nguyen *et al.*

**Supplementary Figures 1 – 8**
**Supplementary Tables 1 – 7**
**Supplementary Data 1**
**Supplementary References**

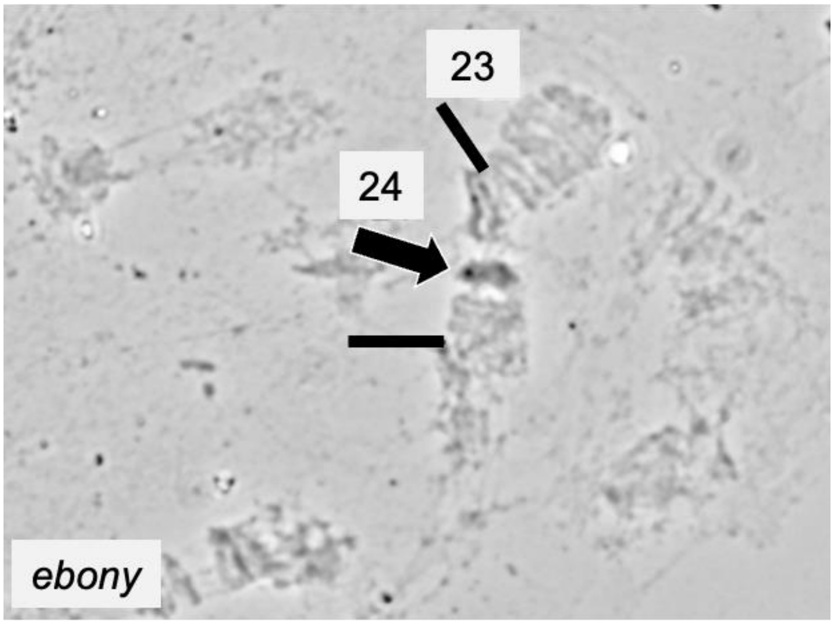


**Supplementary Figure 1.** *In situ* hybridization of *ebony* (arrowhead) in region 24 of the polytene chromosome 3 (mitotic chromosome 2) of *Anastrepha ludens*. Hybridizations were performed in duplicates and at least ten nuclei were analyzed per sample.

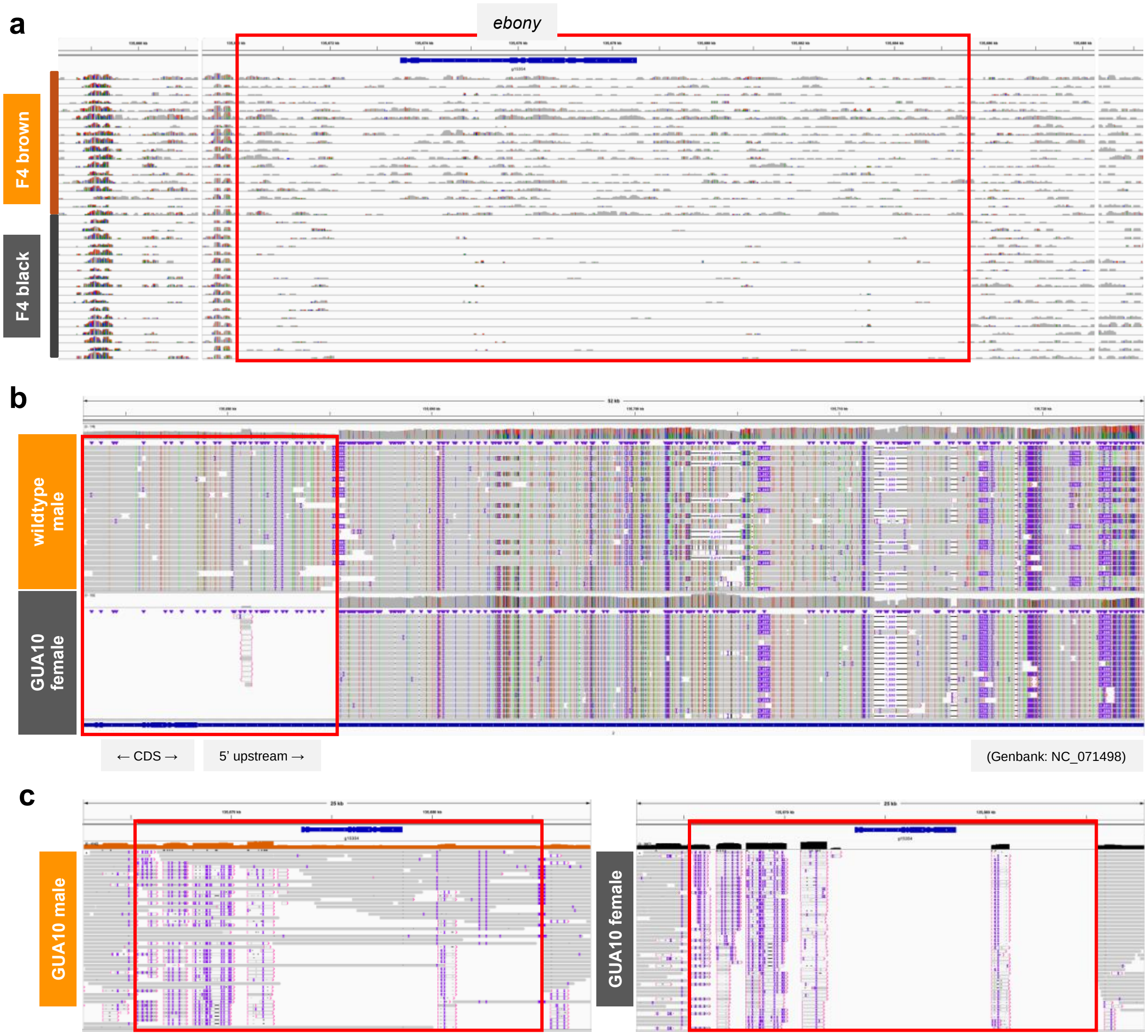


**Supplementary Figure 2.** Localization of the *bp*$^{-}$ mutation within the *ebony* loci of the *A. ludens* GUA10 genetic sexing strain. **(a)** WGS short-read coverage comparison between brown (*bp*$^{+/+}$ or *bp*$^{+/-}$) and black (*bp*$^{-/-}$) pupae siblings from the F4 mapping population (*n* = 18 individuals per phenotype). **(b)** HiFi long-read mapping comparison between wildtype male (*bp*$^{+/+}$) and GUA10 female (*bp*$^{-/-}$). **(c)** HiFi long-read mapping comparison between GUA10 male (*bp*$^{+/-}$) and GUA10 female (*bp*$^{-/-}$). Differential read mapping between samples is highlighted within red boxes. Images are screenshots from the Integrative Genomics Viewer (IGV).

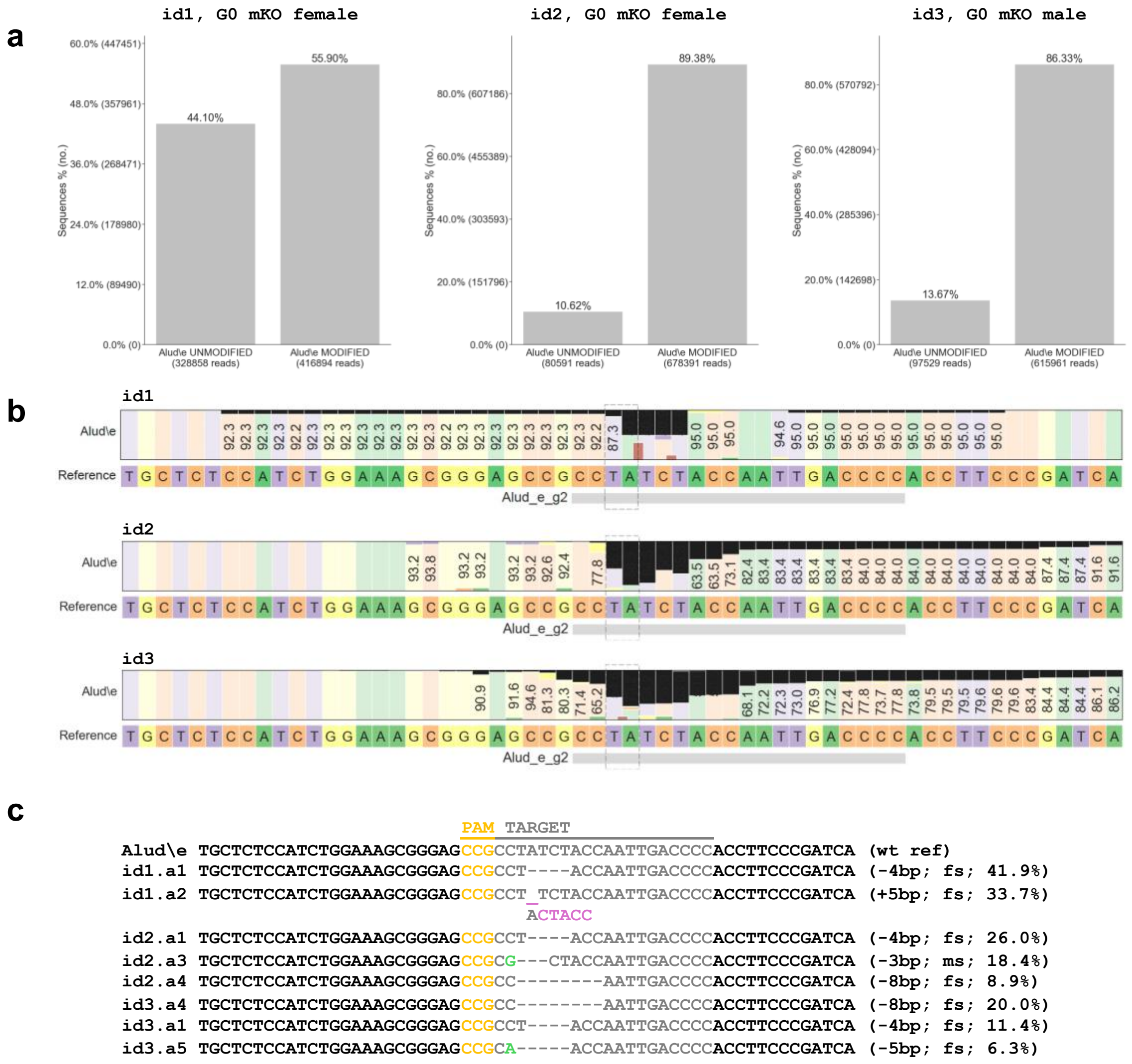


**Supplementary Figure 3.** Illumina genotyping of indexed amplicons surrounding Cas9 cut sites in three individuals (id1–3) of *A. ludens* G0 mosaic knockouts (mKO). The CRISPResso2 pipeline was used to assemble overlapping read pairs and detect indels. **(a)** Mutagenesis frequency as determined by the percentage of reads displaying modified alleles (Quantification window center = -3, Quantification window size = 2 bp). **(b)** Nucleotide distribution across the amplicon and percentage of each base relative to the wildtype reference amplicon (wt ref). Black and brown bars represent the percentage of deletions and insertions, respectively. **(c)** The most frequent allele variants found in mosaic flies. Percentages are relative to 713,490 – 758,982 mapped reads. Expected outcomes include a number of frameshift (fs) and missense (ms) mutations.

**a**

```
                                   PAM TARGET
Afra\e CTCAGTGTCCGAACTGTGGGGGCCGCTGATGTGTGGACTTTCTATTTTAGTTGTACCCAAAGTAATAACTA (wt ref)
        ..S..V..S..E..L..W..G..P..L..M..C..G..L..S..I..L..V..V..P..K..V..I..T..
G2.KO  CTCAGTGTCCGAACTGTGGGGGCCGCTG-------GACTTTCTATTTTAGTTGTACCCAAAGTAATAACTA (-7bp)
        ..S..V..S..E..L..W..G..P..L.-------.D..F..L..F..*..L..Y..P..K..*..*..L.
```

**b**

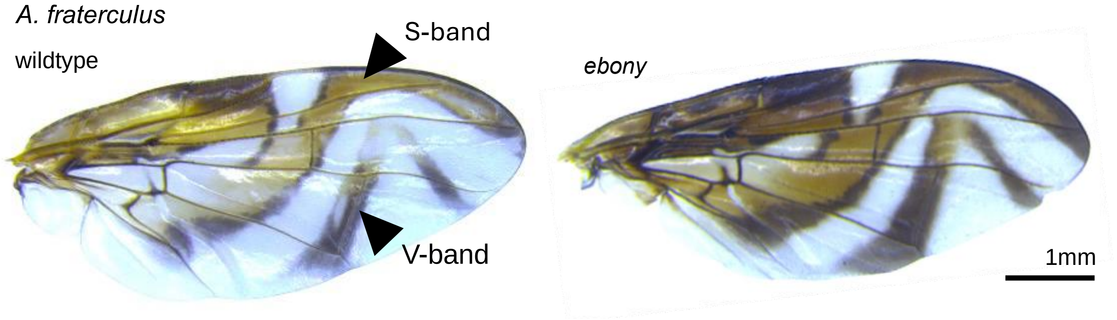


**Supplementary Figure 4. (a)** SANGER sequencing genotyping of *ebony* mutants in *A. fraterculus* G2. Ten out of 36 genotyped flies showed unambiguous results, all displaying a -7 bp deletion resulting in premature stop codons in exon 1 (frameshift mutation). A homozygous mutant line was subsequently established based on this same mutation. **(b)** Disruption of *ebony* leads to alterations in wing pigmentation of *A. fraterculus*, changing S- and V-bands from yellowish-brown to dark-brown with subtle pattern enlargement. No ectopic pigmentation was observed in the wing lamina. Terminology by White and Elson-Harris[3].

**a**

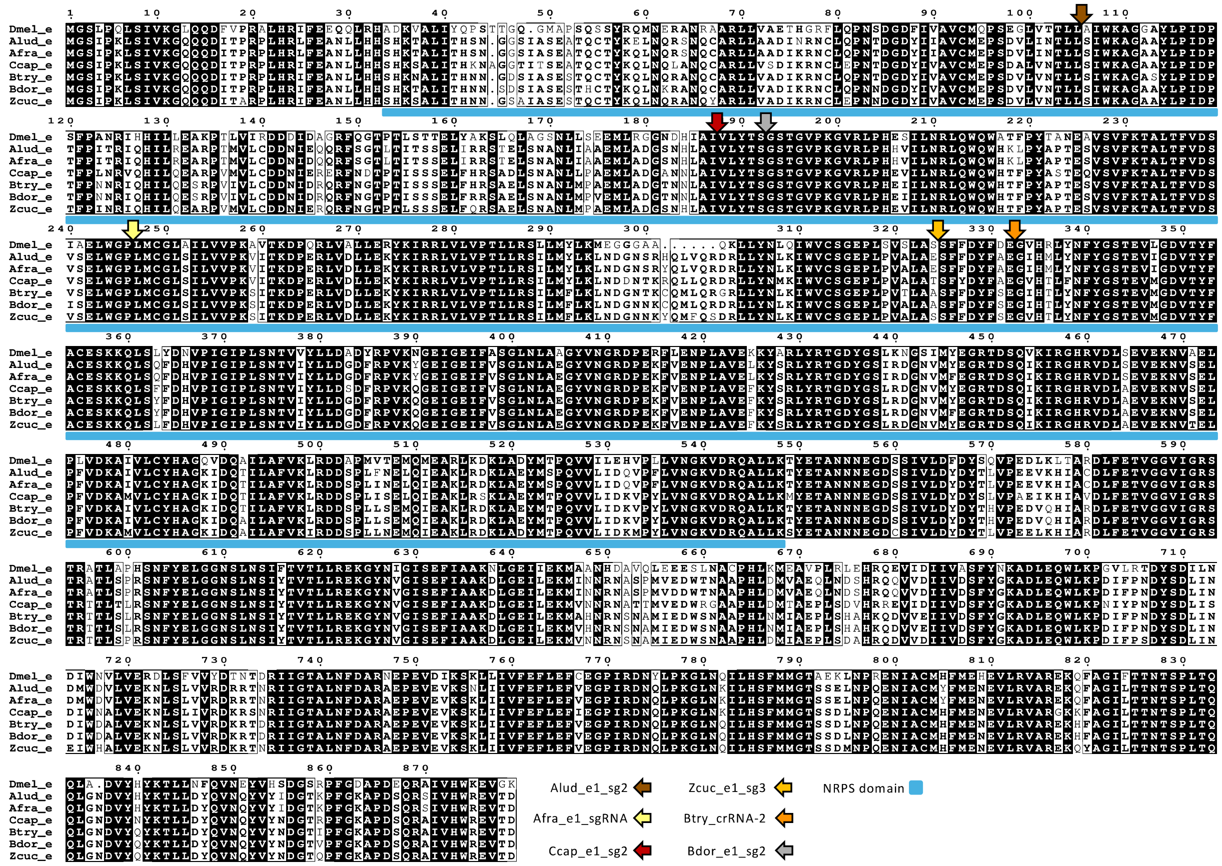


**b**

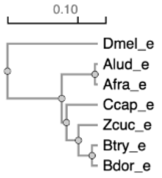


| | | | | | | | |
|---|---|---|---|---|---|---|---|
| 1: Dmel_e | 100.00 | 72.55 | 72.67 | 72.35 | 73.69 | 73.69 | 73.80 |
| 2: Alud_e | 72.55 | 100.00 | 99.10 | 88.37 | 89.50 | 89.16 | 89.50 |
| 3: Afra_e | 72.67 | 99.10 | 100.00 | 88.49 | 89.73 | 89.62 | 89.62 |
| 4: Ccap_e | 72.35 | 88.37 | 88.49 | 100.00 | 90.74 | 90.07 | 90.74 |
| 5: Btry_e | 73.69 | 89.50 | 89.73 | 90.74 | 100.00 | 98.19 | 93.91 |
| 6: Bdor_e | 73.69 | 89.16 | 89.62 | 90.07 | 98.19 | 100.00 | 94.02 |
| 7: Zcuc_e | 73.80 | 89.50 | 89.62 | 90.74 | 93.91 | 94.02 | 100.00 |

**Supplementary Figure 5. (a)** Multiple sequence alignment of Ebony orthologues investigated in this study. Identical residues are shaded in black, while sites with equivalent residues (≥ 70% conservation) are in bold letters inside boxes. Arrowheads indicate approximate sgRNA targeted sites. The non-ribosomal peptide synthetase (NRPS) domain ($S^{34}$ – $K^{548}$), responsible for the Ebony-catalyzed binding of dopamine to β-alanine[1,2], is underlined. **(b)** Phylogenetic relationship between Ebony orthologues (leftmost) and their genetic distance (rightmost). Manually curated gene models (Supplementary Data 1) were translated using the ExPASy, aligned with Clustal Omega, and visualized in ESPript. The Ebony sequence from *Drosophila* (FlyBase: FBgn0000527) was used as an outgroup. Key: Dmel = *Drosophila melanogaster*, Alud = *Anastrepha ludens*, Afra = *Anastrepha fraterculus*, Ccap = *Ceratitis capitata*, Zcuc = *Zeugodacus cucurbitae*, Btry = *Bactrocera tryoni*, and Bdor = *Bactrocera dorsalis*.

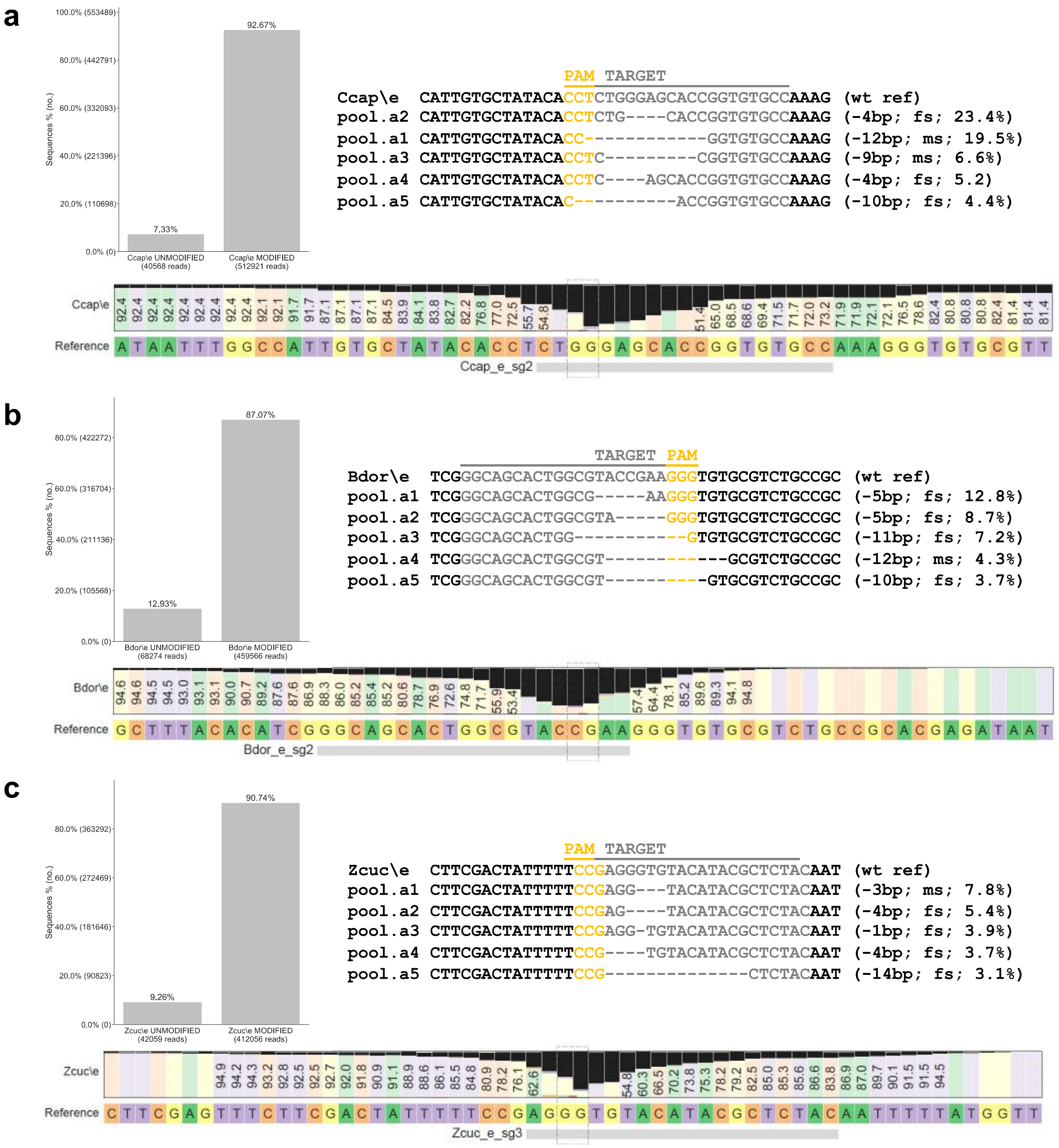


**Supplementary Figure 6.** Illumina genotyping of indexed amplicons surrounding Cas9 cut sites in G0 mosaic knockouts (mKO) of **(a)** *Ceratitis capitata*, **(b)** *Bactrocera dorsalis*, and **(c)** *Zeugodacus cucurbitae*. The CRISPResso2 pipeline was used for analysis (Quantification window center = -3, Quantification window size = 2 bp). Mutagenesis rate (leftmost) is defined as the percentage of reads displaying modified alleles, as shown in the most frequent variants (rightmost). Nucleotide distribution across the amplicon (below) reveals deletions (black bars) and insertions (brown bars) at the targeted sites. Each sample is a pool of mosaic G0 adults (*n* = 3). Percentages are relative to 454,115 – 553,489 mapped reads. Expected outcomes include a number of frameshift (fs) and missense (ms) mutations.

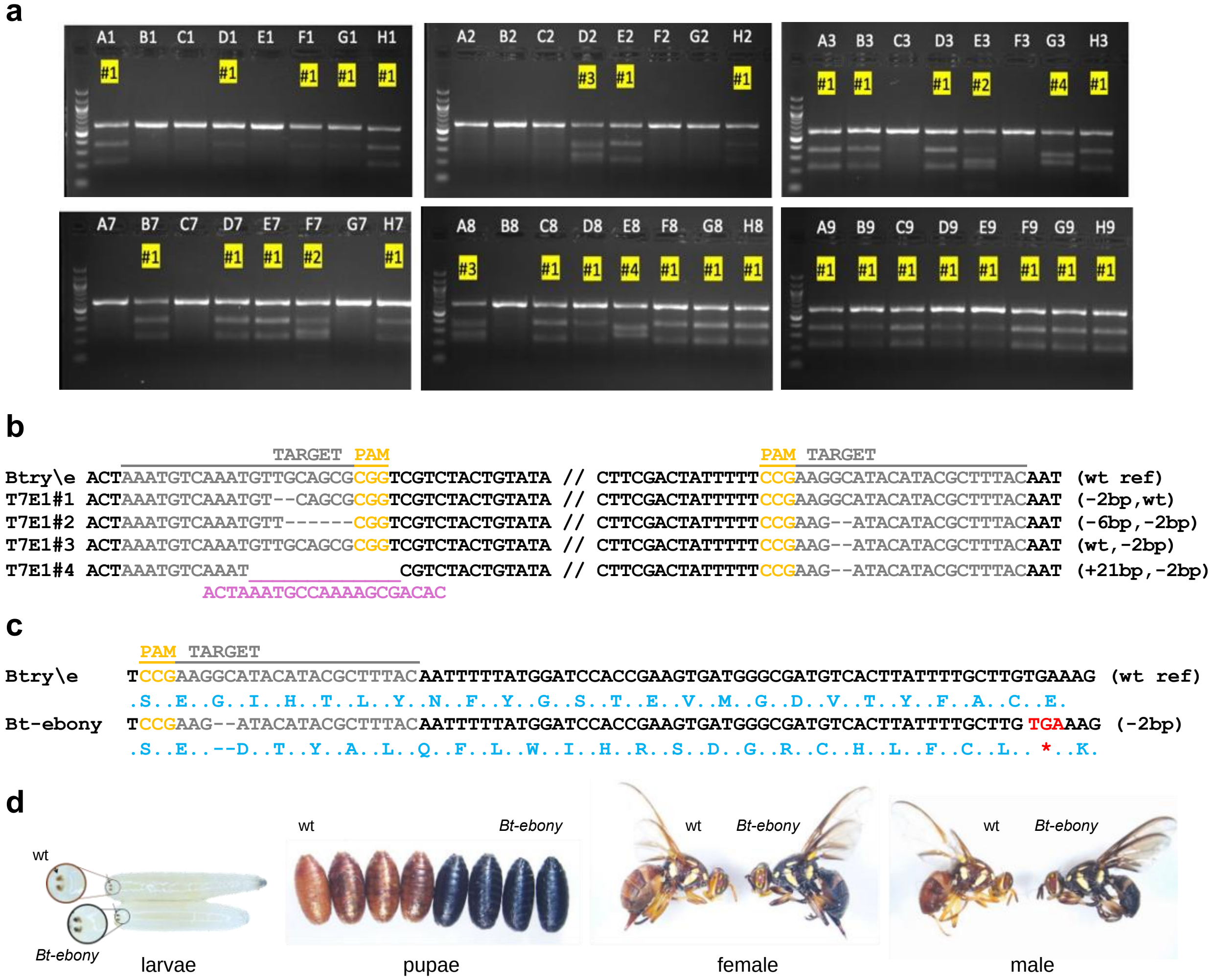


**Supplementary Figure 7.** Genotyping efforts during the establishment of a homozygous *ebony* mutant strain in *Bactrocera tryoni* (*Bt-ebony*). **(a)** T7 endonuclease I (T7EI) assays were used to identify heterozygous mutants from a pool of G2 individuals consisting of 24 males (A1 – H3) and 24 females (A7 – H9). Cleaved products indicate the presence of indels in the sample, and four distinct cleavage patterns were identified (highlighted in yellow). **(b)** SANGER sequencing of positive heterozygous G2 individuals revealed four *ebony* mutant alleles containing indels at one or both Cas9 targeted sites. **(c)** The homozygous mutant line *Bt-ebony* was established based on a -2 bp deletion, leading to premature stop codons in exon 1 (frameshift mutation) and thus *ebony* loss-of-function. **(d)** *ebony* mutants exhibit increased melanin production, resulting in black anal lobes in larvae, black puparium, and darker adult bodies compared to wildtype flies. As in other tephritids studied here, these phenotypes are comparable to the black pupae mutant in *A. ludens*[4].

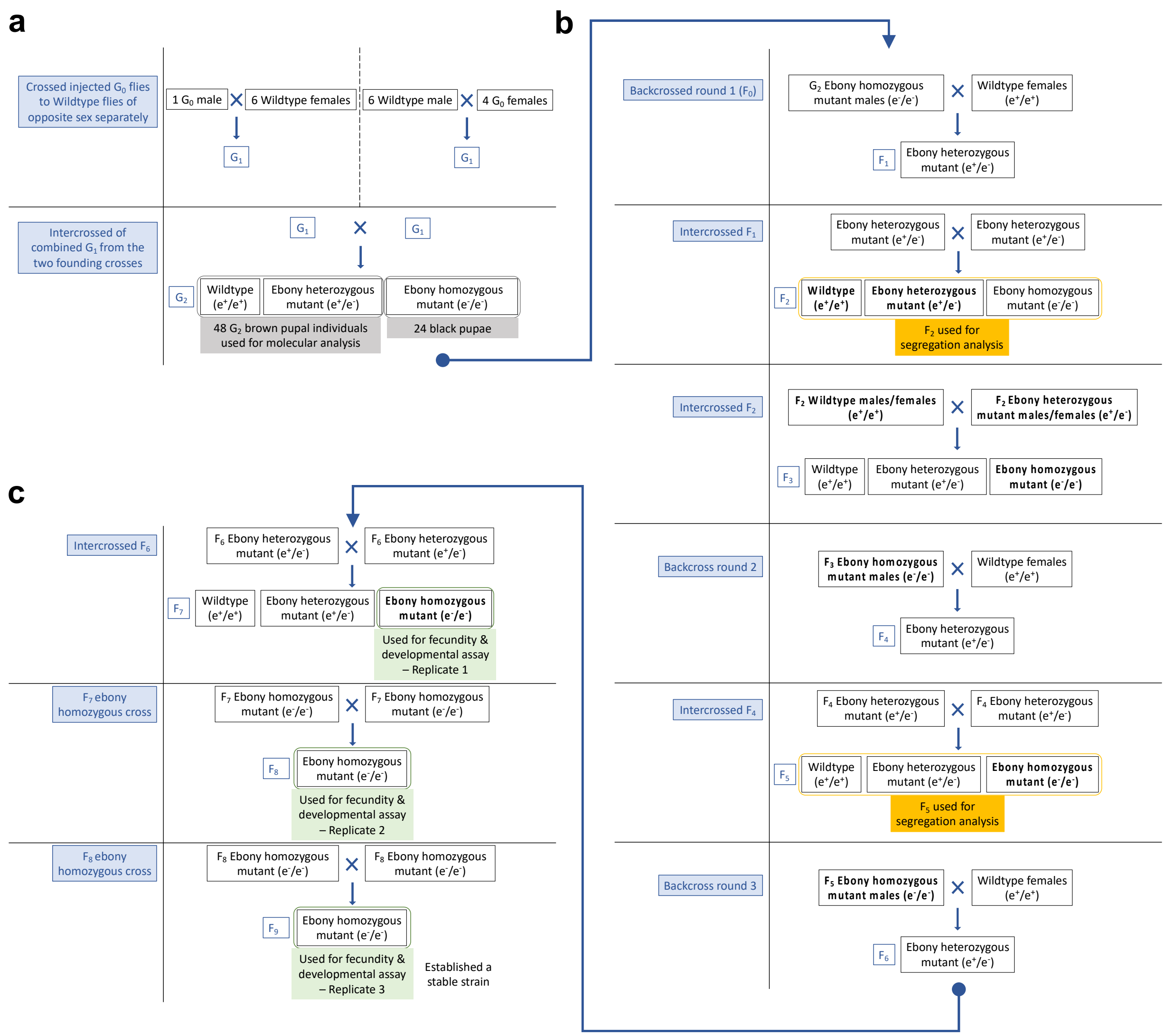


**Supplementary Figure 8.** Crossing scheme leading to a homozygous *ebony* mutant strain in *Bactrocera tryoni* (*Bt-ebony*). **(a)** Following microinjections, surviving G0 flies were backcrossed to their wildtype counterparts. All resulting G1 offspring were mass interbred, and *ebony* mutants recovered at G2. **(b)** To mitigate potential off-target effects from CRISPR/Cas9 experiments, homozygous *ebony* mutants harboring a -2 bp deletion were subjected to three additional rounds of backcrossing to the wildtype strain. Segregation analysis was conducted during the F2 and F5 generations. **(c)** Additional inbreeding was performed between homozygous mutants to establish the *Bt-ebony* strain. Fecundity and developmental assays were conducted with flies from the F7, F8, and F9 generations.

**Supplementary Table 1.** Gene Ontology (GO) terms for Biological Processes enriched in the bp causal region identified in the F4 mapping population. 'Annotated' indicates *A. ludens* genes annotated with the GO term. 'Significant' refers to genes in the causal region with the GO term. 'Expected' is the number of genes within the causal region with the GO term expected by sampling chance. Fisher exact test was used to calculate the *p*-values.

| GO ID | Term | Annotated | Significant | Expected | *p*-value |
|---|---|---|---|---|---|
| GO:0007501 | mesodermal cell fate specification | 45 | 6 | 0.37 | 1.50E-06 |
| GO:0021527 | spinal cord association neuron differentiation | 15 | 3 | 0.12 | 2.20E-04 |
| GO:0051450 | myoblast proliferation | 15 | 3 | 0.12 | 2.20E-04 |
| GO:0007483 | genital disc morphogenesis | 22 | 3 | 0.18 | 7.10E-04 |
| GO:0007218 | neuropeptide signaling pathway | 88 | 5 | 0.72 | 7.20E-04 |
| GO:0048665 | neuron fate specification | 51 | 4 | 0.41 | 7.60E-04 |
| GO:0008340 | determination of adult lifespan | 323 | 9 | 2.63 | 1.17E-03 |
| GO:0001709 | cell fate determination | 210 | 7 | 1.71 | 1.54E-03 |
| GO:0010259 | multicellular organism aging | 343 | 9 | 2.79 | 1.78E-03 |
| GO:0021522 | spinal cord motor neuron differentiation | 31 | 3 | 0.25 | 1.97E-03 |
| GO:0030218 | erythrocyte differentiation | 69 | 4 | 0.56 | 2.34E-03 |
| GO:0034243 | regulation of transcription elongation by RNA polymerase II | 35 | 3 | 0.28 | 2.81E-03 |
| GO:0090110 | COPII-coated vesicle cargo loading | 10 | 2 | 0.08 | 2.82E-03 |
| GO:0061320 | pericardial nephrocyte differentiation | 10 | 2 | 0.08 | 2.82E-03 |
| GO:0017148 | negative regulation of translation | 178 | 6 | 1.45 | 3.19E-03 |
| **GO:0006583** | **melanin biosynthetic process from tyrosine** | **11** | **2** | **0.09** | **3.42E-03** |
| GO:0048047 | mating behavior, sex discrimination | 11 | 2 | 0.09 | 3.42E-03 |
| GO:0009408 | response to heat | 183 | 6 | 1.49 | 3.65E-03 |
| GO:0042684 | cardioblast cell fate commitment | 12 | 2 | 0.1 | 4.09E-03 |
| GO:0007368 | determination of left/right symmetry | 133 | 5 | 1.08 | 4.46E-03 |
| GO:0042659 | regulation of cell fate specification | 83 | 4 | 0.67 | 4.57E-03 |
| GO:0045893 | positive regulation of DNA-templated transcription | 896 | 15 | 7.29 | 5.11E-03 |
| GO:0019184 | nonribosomal peptide biosynthetic process | 14 | 2 | 0.11 | 5.57E-03 |
| GO:0007480 | imaginal disc-derived leg morphogenesis | 90 | 4 | 0.73 | 6.09E-03 |
| GO:0071236 | cellular response to antibiotic | 145 | 5 | 1.18 | 6.42E-03 |
| GO:0031327 | negative regulation of cellular biosynthetic process | 1172 | 21 | 9.53 | 7.21E-03 |
| GO:0010628 | positive regulation of gene expression | 1321 | 19 | 10.74 | 8.20E-03 |
| GO:0061418 | obsolete regulation of transcription from RNA polymerase II promoter in response to hypoxia | 17 | 2 | 0.14 | 8.20E-03 |
| GO:1904385 | cellular response to angiotensin | 17 | 2 | 0.14 | 8.20E-03 |
| GO:0045451 | pole plasm oskar mRNA localization | 53 | 3 | 0.43 | 9.05E-03 |
| GO:0002474 | antigen processing and presentation of peptide antigen via MHC class I | 18 | 2 | 0.15 | 9.18E-03 |
| GO:1904037 | positive regulation of epithelial cell apoptotic process | 18 | 2 | 0.15 | 9.18E-03 |
| GO:0035050 | embryonic heart tube development | 102 | 4 | 0.83 | 9.41E-03 |
| GO:0009064 | glutamine family amino acid metabolic process | 54 | 3 | 0.44 | 9.52E-03 |
| GO:0051241 | negative regulation of multicellular organismal process | 781 | 13 | 6.35 | 9.71E-03 |

**Supplementary Table 2.** Differentially expressed genes (DEGs) between 1d-old black (*n* = 3, females) and brown (*n* = 3, males) pupae siblings of *A*. *ludens* GUA10 strain identified by edgeR. Overall, expression differences among 10,913 genes were not significant, with only four genes found to be upregulated in brown pupae samples. logFC = log2 fold-change between tested conditions (*-1*black 1*brown*), FDR = false discovery rate.

| Feature | logFC | *p*-value | FDR | Gene | Chr. (RefSeq) | Function (FlyBase) |
|---|---|---|---|---|---|---|
| g794 | 8.2142 | 3.74E-07 | 0.0041 | *Arc1* | 6 (NC_071502.1) | Master regulator of synaptic plasticity (FBgn0033926). |
| g15354 | 7.4408 | 8.29E-06 | 0.0302 | *ebony* | 2 (NC_071498.1) | Links beta-alanine to biogenic amines like dopamine. Involved in cuticle pigmentation (FBgn0000527) |
| g13426 | 4.7262 | 1.16E-05 | 0.0316 | unknown | 2 (NC_071498.1) | Unknown. Also found in *Ceratitis* and *Bactrocera* (BLASTp). |
| g15461 | 1.6355 | 8.10E-06 | 0.0302 | *Spc25* | 2 (NC_071498.1) | Component of the Ndc80 complex, which is an essential kinetochore constituent (FBgn0087021) |

**Supplementary Table 3.** Summary statistics of CRISPR/Cas9 experiments leading to the disruption of *ebony* in diverse tephritids. mKO = mosaic knockout, wt = wild-type.

| Species | Eggs | Surviving G0 pupae | | | Crossing | Offspring (G1 or G2) | | |
|---|---|---|---|---|---|---|---|---|
| | | Total | Black | Brown | | Total | Black | Brown |
| *A. ludens* | 260 | 38 (14.6%) | 4 (10.5%) | 34 (89.5%) | mass G0 x G0 | no data | none | all (100%) |
| *A. fraterculus* | 179 | 24 (13.4%) | none | all (100%) | ind. G0 x wt → mass G1 x G1 | no data | 140 | no data |
| *C. capitata* | 260 | 69 (26.5%) | 40 (58.0%) | 29 (42.0%) | mass mKO x mKO | 247 | 194 (78.5%) | 53 (21.5%) |
| *B. triony* | 661 | 12 (1.8%) | none | all (100%) | group G0 x wt → mass G1 x G1 | no data | 24 | no data |
| *B. dorsalis* | 240 | 82 (34.2%) | 56 (68.3%) | 26 (31.7%) | mass mKO x mKO | 266 | 244 (91.7%) | 22 (8.3%) |
| *Z. cucurbitae* | 252 | 38 (15.1%) | 26 (68.4%) | 12 (31.6%) | mass mKO x mKO | 441 | 266 (60.3%) | 175 (39.7%) |

**Supplementary Table 4.** Phenotype and genotype segregation analyses of *ebony* in *B. tryoni*. **(a)** F2 progeny from crosses between *B. tryoni ebony* mutants and wildtype flies exhibit no deviation from the expected 3:1 phenotype and 1:2 genotype ratios. Gender distribution based on F2 phenotypes **(b)** and wildtype brown pupae genotypes **(c)** in replicate 2 shows no bias in the expected 1:1 sex ratio.

**a**

| **F0 parents** | | **Rep.** | **F2 phenotypes** | | **Chi-square against 3:1 ratio** | | | **F2 brown genotypes** | | **Chi-square against 1:2 ratio** | | |
|---|---|---|---|---|---|---|---|---|---|---|---|---|
| **Male** | **Female** | | **brown** | **black** | **X2** | **df** | ***p*** | **e +/+** | **e +/-** | **X2** | **df** | ***p*-value** |
| *ebony* | wildtype | #1 | 43 | 18 | 0.661 | 1 | 0.416 | 10 | 22 | 0.062 | 1 | 0.802 |
| *ebony* | wildtype | #2 | 157 | 42 | 1.609 | 1 | 0.204 | 15 | 33 | 0.093 | 1 | 0.759 |
| ***Overall stats*** | | | **200** | **60** | **0.513** | **1** | **0.474** | **25** | **55** | **0.156** | **1** | **0.692** |

**b**

| **F2 phenotypes** | **Sex** | | | **Chi-square against 1:1 ratio** | | |
|---|---|---|---|---|---|---|
| **Rep. #2** | **Male** | **Female** | **Total** | **X2** | **df** | ***p*-value** |
| brown | 78 | 79 | 157 | 0.006 | 1 | 0.936 |
| black | 19 | 23 | 42 | 0.380 | 1 | 0.537 |
| ***Overall stats*** | **97** | **102** | **199** | **0.125** | **1** | **0.723** |

**c**

| **F2 brown genotypes** | **Sex** | | | **Chi-square against 1:1 ratio** | | |
|---|---|---|---|---|---|---|
| **Rep. #2** | **Male** | **Female** | **Total** | **X2** | **df** | ***p*-value** |
| *ebony +/+* | 8 | 7 | 15 | 0.666 | 1 | 0.796 |
| *ebony +/-* | 16 | 17 | 33 | 0.030 | 1 | 0.861 |
| ***Overall stats*** | **24** | **24** | **48** | **0.000** | **1** | **1.000** |

**Supplementary Table 5.** Source data for fitness analysis of *Bt-ebony* KO mutants. The experiment included four genetic crossing combinations (control, test, reciprocal, and ebony), each performed with three independent replicates. Each cross was composed of 10 males and 10 females. Data recorded at the F1 generation included counts of eggs, larvae, pupae, partially emerged adults, fully emerged adults, normal emerged adults, and deformed emerged adults.

| Crossing | Exp. | Rep. | Phenotype | Eggs | Larvae | Pupae | Partial | Full | Normal | Deformed |
|---|---|---|---|---|---|---|---|---|---|---|
| wt(M)_vs_wt(F) | control | 1 | brown | 2399 | 1925 | 1325 | 7 | 1116 | 1109 | 7 |
| wt(M)_vs_wt(F) | control | 2 | brown | 927 | 617 | 451 | 4 | 425 | 419 | 6 |
| wt(M)_vs_wt(F) | control | 3 | brown | 3327 | 2829 | 2229 | 12 | 1793 | 1780 | 13 |
| ebony(M)_vs_wt(F) | test | 1 | brown | 1359 | 903 | 661 | 2 | 562 | 552 | 10 |
| ebony(M)_vs_wt(F) | test | 2 | brown | 376 | 88 | 37 | 1 | 36 | 35 | 1 |
| ebony(M)_vs_wt(F) | test | 3 | brown | 3649 | 2461 | 1423 | 5 | 1208 | 1195 | 13 |
| wt(M)_vs_ebony(F) | reciprocal | 1 | brown | 1992 | 1337 | 907 | 5 | 825 | 807 | 18 |
| wt(M)_vs_ebony(F) | reciprocal | 2 | brown | 720 | 457 | 270 | 1 | 260 | 255 | 5 |
| wt(M)_vs_ebony(F) | reciprocal | 3 | brown | 1847 | 999 | 685 | 2 | 627 | 618 | 9 |
| ebony(M)_vs_ebony(F) | ebony | 1 | black | 1469 | 535 | 363 | 24 | 296 | 282 | 14 |
| ebony(M)_vs_ebony(F) | ebony | 2 | black | 744 | 225 | 163 | 44 | 103 | 96 | 7 |
| ebony(M)_vs_ebony(F) | ebony | 3 | black | 1355 | 651 | 352 | 32 | 248 | 241 | 7 |

**Supplementary Table 6.** List of sequencing data from *A. ludens* used in this study. Dataset included sequences generated by Gutiérrez-Ramos et al[5]. and Sirot et al.[6].

| SRA accession | Data | Sample | Purpose | Reference |
|---|---|---|---|---|
| SRR11028485 | RNA-Seq | wildtype embryos 0-9h AEL | structural genome annotation | Gutiérrez-Ramos et al. |
| SRR11028488 | RNA-Seq | wildtype embryos 9-18h AEL | structural genome annotation | Gutiérrez-Ramos et al. |
| SRR11028491 | RNA-Seq | wildtype embryos 18-30h AEL | structural genome annotation | Gutiérrez-Ramos et al. |
| SRR8612579 | RNA-Seq | GUA10 female pupa | structural genome annotation | *unpublished data* |
| SRR8612578 | RNA-Seq | GUA10 male pupa | structural genome annotation | *unpublished data* |
| SRR8612576 | RNA-Seq | wildtype virgin female | structural genome annotation | *unpublished data* |
| SRR8612577 | RNA-Seq | wildtype mated female | structural genome annotation | *unpublished data* |
| SRR9841864 | RNA-Seq | wildtype naive male | structural genome annotation | Sirot et al. |
| SRR9841863 | RNA-Seq | wildtype mated male | structural genome annotation | Sirot et al. |
| SRR0000000 - 0000000 | WGS | F4 mapping population, brown pupae | identification of the bp causal region | this study |
| SRR0000000 - 0000000 | WGS | F4 mapping population, black pupae | identification of the bp causal region | this study |
| SRR0000000 - 0000000 | RNA-Seq | GUA10 female pupa | DGE analysis | this study |
| SRR0000000 - 0000000 | RNA-Seq | GUA10 male pupa | DGE analysis | this study |
| SRR17880705 | HiFi-Seq | wildtype (Willacy) adult male | characterization of the *bp* mutation | *unpublished data* |
| SRR0000000 | HiFi-Seq | GUA10 adult female | characterization of the *bp* mutation | this study |
| SRR0000000 | HiFi-Seq | GUA10 adult male | characterization of the *bp* mutation | this study |

**Supplementary Table 7.** List of primers used in this study.

| **Primer name** | **Sequence (5' → 3')** | **Purpose** |
|---|---|---|
| Alud_e_F1 | GCCAGTTCGATCATGTACCG | PCR (360bp) and RT-PCR (188bp) spanning Alud\e exons e1 and e2 |
| Alud_e_R2 | TTCCACAGCCAATGGATTTTCG | |
| Alud_RpL18_F1 | AACTGAGCCCAAATCGCAAG | PCR (351bp) and RT-PCR (156bp) spanning Alud\RpL18 exons e1 and e2 |
| Alud_RpL18_R1 | CTGACACGCTGCAAAGACAT | |
| Alud_ebony_probe_new_F | CTGGCTCAATTGCTTTTTGCT | PCR generated *in situ* probe for Alud\e |
| Alud_ebony_probe_new_R | AACTGCACTGATAACGCACAG | |
| Alud_e1_sg2 | GAAATTAATACGACTCACTATAGGGGTCAATTGGTAGATAGGGTTTTAGAGCTAGAAATAGC | synthesis of sgRNA against Alud\e |
| Alud_e1_sg2_fwd.p5 | TCGTCGGCAGCGTCAGATGTGTATAAGAGACAGTGTGCTCATCAATACGCTGC | Illumina genotyping of PCR amplicons (173bp) spanning guide recognition site in Alud\e |
| Alud_e1_sg2_rev.p7 | GTCTCGTGGGCTCGGAGATGTGTATAAGAGACAGTAATGGTGAGTGTGCCGCTA | |
| Afra_e1_sgRNA | ATAGAAAGTCCACACATCAG | custom sgRNA against Afra\e |
| GMB_174_rev | ACATCGCCTATCACTTCGGT | SANGER genotyping of PCR amplicons spanning guide recognition site in Afra\e |
| GMB_179_fwd | GCACAACAATGGCGGTTCAA | |
| Ccap_e1_sg2 | GAAATTAATACGACTCACTATAGGCACACCGGTGCTCCCAGGTTTTAGAGCTAGAAATAGC | synthesis of sgRNA against Ccap\e. |
| Ccap_e1_sg2_fwd.p5 | TCGTCGGCAGCGTCAGATGTGTATAAGAGACAGCAGCAGCTCGGAACTCTTTC | Illumina genotyping of PCR amplicons (211bp) spanning guide recognition site in Ccap\e |
| Ccap_e1_sg2_rev.p7 | GTCTCGTGGGCTCGGAGATGTGTATAAGAGACAGCTCTGTGGACGCATAAGGGA | |
| Btry_crRNA-1 | AAATGTCAAATGTTGCAGCG | custom crRNA against Btry\e |
| Btry_crRNA-2 | GTAAAGCGTATGTATGCCTT | custom crRNA against Btry\e |
| Qebony_F1 | CAAGACCGCACTCACCTTTG | T7E1 and SANGER genotyping of PCR amplicons spanning guide recognition sites in Btry\e |
| Qebony_R1 | CTCCTTGTTTGACTGGTCGG | |
| Bdor_e1_sg2 | GAAATTAATACGACTCACTATAGGCAGCACTGGCGTACCGAAGTTTTAGAGCTAGAAATAGC | synthesis of sgRNA against Bdor\e |
| Bdor_e1_sg2_fwd.p5 | TCGTCGGCAGCGTCAGATGTGTATAAGAGACAGACGCTAATTTGATGCCGGYC | Illumina genotyping of PCR amplicons (198bp) spanning guide recognition site in Bdor\e |
| Bdor_e1_sg2_rev.p7 | GTCTCGTGGGCTCGGAGATGTGTATAAGAGACAGTGAGTGCGGTCTTGAAAACG | |
| Zcuc_e1_sg3 | GAAATTAATACGACTCACTATAGGTAGAGCGTATGTACACCCTGTTTTAGAGCTAGAAATAGC | synthesis of sgRNA against Zcuc\e |
| Zcuc_e1_sg3_fwd.p5 | TCGTCGGCAGCGTCAGATGTGTATAAGAGACAGCCGTTTAGTTTTGGTGCCGA | Illumina genotyping of PCR amplicons (237bp) spanning guide recognition site in Zcuc\e |
| Zcuc_e1_sg3_rev.p7 | GTCTCGTGGGCTCGGAGATGTGTATAAGAGACAGACGTCACCCATTACTTCRGT | |

**Supplementary Data 1.** Manually curated annotation of *ebony* orthologues investigated in this study (in FASTA format). Exons are in uppercase. Introns are in lowercase within curly brackets.

```
>Anastrepha_ludens_ebony
ATGGGATCTATACCAAAACTATCAATTGTTAAAGGCCAACAACAGGATATAACGCCCCGTCCGT
TGCATCGCCTTTTTGAGGCAAATCTGTTGCATCACTCGCACAAAACTGCACTGATAACGCACAG
CAATGGCGGTTCAATCGCATCTGAAGCCACGCAATGCACTTACAAGGAATTGAATCAGCGCTCC
AACCAATGTGCACGTCTACTAGCGGCTGACATAAACCGTAACTGTTTGCAACCTAACACCGACG
GCGATTATATTATAGCAGTATGCATGGAACCATCAGATGTGCTCATCAATACGCTGCTCTCCAT
CTGGAAAGCGGGAGCCGCCTATCTACCAATTGACCCCACCTTCCCGATCACACGCATACAACAT
ATCCTACGAGAAGCGCGTCCGACTATGGTGCTATGTGATGACAATATTGAACAACAACGCTTTA
GCGGCACACTCACCATTACAAGCTCGGAACTCATTCGCCGTTCCACCGAGTTAAGTAACGCAAA
CTTAATTGCTGCCGAGATGCTGGCCGACGGTAGTAATCATTTGGCGATTGTGCTCTACACATCG
GGCAGTACTGGTGTACCAAAAGGCGTGCGCCTACCACATCACGTAATCTTGAACCGTCTGCAAT
GGCAATGGCACAAATTGCCTTACGCGCCAACTGAGTCGGTGAGCGTTTTCAAGACAGCACTCAC
ATTCGTCGACTCAGTGTCCGAACTGTGGGGGCCGCTGATGTGTGGACTTTCTATTTTAGTTGTA
CCCAAAGTAATAACTAAAGATCCGGAGCGTTTGGTCGATTTGCTAGAGAAATATAAAATACGTC
GTTTAGTTTTGGTGCCAACACTTTTGCGTTCTATTTTGATGTATTTGAAGTTGAACGATGGCAA
TTCAAGACATCAACTGGTGCAACGCGACCGACTTTTGTATAATTTGAAGATTTGGGTTTGTTCT
GGTGAACCTTTGCCGGTAGCACTGGCAGAAAGTTTCTTTGACTACTTCGCTGAGGGCATACATA
TGCTGTATAACTTTTACGGTTCCACCGAAGTGATAGGTGATGTTACGTACTTTGCTTGTGAAAG
CAAAAAGCAATTGAGCCAGTTCGATCATGTACCGATTG{gtgagtcgaaactttgagatattca
agcttttccaataaacgtcttaatataaatttaactgctggtttaattcataataacagcttat
ttcatagcatgggcggcgattagtgtcgtgatctgtttgtacttaacacttttctaacagttta
cgttttcacccgtgatag}GCATTCCACTTTCCAACACTGTTATCTACTTACTTGATGGAGACT
TCCGCCCAGTCAAATATGGCGAAATAGGTGAAATATTTGTTTCGGGCCTGAATCTTGCCGAAGG
TTATGTAAATGGTAGAGACCCGGAAAAGTTTGTCGAAAATCCATTGGCTGTGGAATTAA{gtaa
gttcaaacaaatcatccctctatgtaagtacgtaaatgaatacaattcctcttcgtcacaaaca
g}AATACTCACGTCTATATCGCACTGGTGATTATGGCTCTATACGGGATGGCAATGTGATGTAC
GAAGGCCGAACCGATTCGCAAATAAAGATTCGTGGTCATCGTGTCGATCTTAGCGAAGTTGAAA
AGAACGTTTCAGAGTTGCCGTTCGTCGATAAAGCGATCGTACTTTGCTATCATGCCGGCAAAAT
TGATCAGACGATTCTAGCTTTTGTTAAGCTACGCGACGATTCACCGCTATTCAACGAACTACAA
ATCGAGGCTAAATTGAGAGATAAGCTCGCGGAGTACATGTCACCACAAGTTGTGCTTATTGACC
AAGTTCCATTCCTTGTCAATGGGAAAGTAGATCGGCAGGCACTTCTCAAAACTTACGAGACAGC
CAATAATAACGAAGGCGATTCCAGTATTGTGCTGGACTATGATTATACACTCGTACCCGAAGAG
GTAAAGCACATTGCCTGTGATCTCTTTGAGACCGTGGGTGGCGTAATCGGTCGTTCTACACGCG
CCACTCTATCGCCTCGCAGCAACTTTTACGAGTTGGGCGGAAATTCGTTGAATTCGATTTACAC
AGTCACACTGTTGAGAGAAAAGGGTTACAATGTAGGCATATCTGAATTTATTGCCGCAAAGGAT
TTGGGTGAAATATTGGAGAAAATGATAAATAATCGCAATGCCTCCCCAATGGTCGAGGACTGGA
CAAACGCTGCGCCACATCTTGATATGGTGGCTGAGCAGCTCAATGATTCCCACAGGCAGCAAGT
```

GGTGGA{gtaagtaaagtgttgaatgcattaattgcgtatggacatctatattaatgtaattct
cttttttacaaag}CATCATAGTTGACAGCTTTTATGGCAAGGCGGATCTGGAACAATGGCTCA
AACCAGATATATTCCCGAATGACTACAGTGATCTTATTAAT{gtaatatttgatttaattcctt
ccacttatgtatttataagtgaatttgttcttttccttgctatag}GATATGTGGGATGTTTTA
GTAGAGAAGAATCTCAGTTTGGTGGTACGTGACAGACGTACAAATCGCATCATTGGAACGGCCT
TAAATTTTGACGCACGCGCTGAACCCGAAGTTGAGGTTAAGTCAAATCTTATTATAGTATTTGA
GTTTTTAGAGTTTGTTGAAGGCCCCATTCG{gtaagtttactaattatatataggcatatgtat
atatactacataaacttgtacacattcatacgcataagtacgtacatatgtacagggtgatcaa
tttagaggtatcggattttaaattgaaataaaacaacgaaatttcaaattgattaggcaatctt
tattatgtttgtgtagaaccgtagaaccatattctttcaaatggtagccgcaactgcgccttaa
tccgtaaacaccaattttgaaagactcgttggggcacttcgatcggtatctcgtgaataacttt
agtaatgttggcttccaattccccagccgaatctggtttatccacaaagcatttagactttaca
tgcccctacaaattaaagttcaaaattgtgatataacacgattttggtagccaattcactagtc
cgataagagatataaattgttcactgaaacgacatcgccgtgaatctattgtttcacgggctgt
atggcaagtagcgccgtctgccgtcttgttggaacgggcttcaatttccggcaccacggcatca
aaaagtagtttatcatggcgcgatagcgttcgccactcactgttacattgccgccagactcgtc
tttgaagaaatataggtccatgatttctctagcgcataagtcgcaccaatggatgtaatggctt
ttcttgaatggcttcaagttgctcttcagcccaaatatggcaaatttgcttattgacgtagcca
taaagccagaaatgggcctcatcgctactcaccatgcgttggcgtgagcgcgcaatgaacaatt
atcaaagagcgtcgattttcgtaatacaattgtacgattggtaaacgttgttgagaagcaagtc
tttccatgatgaaatgtcatgaattctgaataaaatgatgtatttagtttgacagtagtgaagc
gtgatctgtcaaaaaagccctattgaaaaaagtacctccaatctggtcaccctttataatgttt
tgattgttttgaaaatgctttctgaacgtgattttattgggtacaaatatatagtatcaggtgg
cgcaaaattagtcatcgcgttgaaagacttataattttgcaaatggcgtcgtagatcaatcat
atttgacacttgaaagctatactgctgtagtaaacaaacagacaaactatgaagcgcgtaaaat
tcaaaataaagatttccactcaaaaccatttaataaatcaaatgtattttaaagagttgtttgc
tgttgttgttatttaaaaaattattcaaaaaaaatttatgattaattttgcgccaccttttct
ctactcaagtaactaattcaatattgtcaagttagcatatctgaggaaggcatactagaaatcc
agcttaatttttttacccactaaacgtataccatttcgagtacgtaggttcgaatctccgtgcat
gaaactccaaaatgaagaaaaagtttcttctaatagcggtcgcccctcggcatgcaatggcgaa
gctccgagtatatttctgccttgaaaaagcttctcattccatgtgtggaacaacatcaagacgc
acaccacaaataggagcaggagcttggccaaacaccttacagaagtgtgcgcgcaaattattat
tattttttcttaaacgtatgccatcaatcgagggaagctccaagtagctcattgcatgcgcgct
gaaccctattcatccaaaacacactacgactatatatcgaatgatcttaaaaattagaaattaa
agaattatttaatggagggacaaaaacaaaaaaagttgtatacaaattattcatttttattta
tttataatatttgtccaaactaaaagtctatgcaaaactcagtttgcagttttataattgtttg
catttttttgctttaatttattttaaaataaag}GGACAACCAACTTCCCAAGGGGCTCAACAAA
ATTCTCCACTCGTTCATGATGGGCACCAGCTCCGAATTGAATCCACAAGAGAATATCGCCTGCA
TGTATTTCATGGAAAACGAAGTGCTGCGCGTTGCTAGAGAGAAACAATTTGCCGGAATTTTAAC
AACCAATACAAGTCCCTTAACGCAG{gtaaaacaaaggaaccaaatggttcaaggttttgcaat
atgagcgatttcttatacttttctccgtacaattacag}CAACTCGGTAACGATGTCTACCACT

ACAAGACATTGCTGGACTATCAGGTCAATCAATACGTATACATCGATGGCACTAAACCATTCGG
AAAAGCTCCCGATTCACAACGTTCCATTGTTCATTGGCGCGAAGTTACCGACTAA

>Anastrepha_fraterculus_ebony
ATGGGATCTATACCAAAGCTATCAATTGTTAAAGGCCAACAACAGGATATAACGCCCCGTCCGT
TGCATCGCCTTTTTGAGGCAAATCTGTTGCATCACTCGCACAAAACTGCACTGATAACGCACAG
CAATGGCGGTTCAATCGCATCTGAAGCCACGCAATGCACTTACAAACAACTGAATCAGCGCTCC
AACCAATGTGCACGTCTACTAGCGGCTGACATAAAGCGTAACTGTTTGCAACCTAACACCGACG
GCGATTATATTATAGCAGTATGCATGGAGCCATCAGATGTTCTCATCAATACGCTGCTCTCCAT
CTGGAAAGCGGGAGCCGCCTATCTACCAATTGACCCCACCTTCCCGATCACACGCATACAACAT
ATCCTACGAGAAGCGCGTCCGACTATGGTGCTATGTGATGACAATATTGATCAACAACGCTTTA
GCGGCACACTCACCATTACAAGCTCGGAGCTCATTCGCCGTTCCACCGAGTTAAGTAACGCAAA
CTTAATTGCTGCCGAGATGCTGGCCGACGGTAGTAATCATTTGGCGATTGTGCTCTACACATCG
GGCAGTACTGGTGTGCCAAAAGGCGTGCGCCTACCACATCACGTAATCTTGAACCGTCTGCAAT
GGCAATGGCACAAATTGCCTTACGCGCCAACTGAGTCTGTGAGCGTTTTCAAGACAGCACTCAC
ATTCGTCGACTCAGTGTCCGAACTGTGGGGGCCGCTGATGTGTGGACTTTCTATTTTAGTTGTA
CCCAAAGTAATAACTAAAGATCCGGAGCGTTTGGTCGATTTGCTAGAGAAATATAAAATACGTC
GTTTAGTTTTGGTGCCAACACTTTTGCGTTCGATTTTGATGTATTTGAAGTTGAACGATGGCAA
TTCAAGATATCAACTGGTGCAACGCGACCGACTTTTGTATAATTTGAAGATTTGGGTTTGTTCT
GGTGAACCTTTGCCGGTAGCACTGGCAGAAAGTTTCTTTGACTACTTCGCTGAGGGCATACATA
TGCTGTATAACTTTTACGGTTCCACCGAAGTGATAGGCGATGTTACGTACTTTGCTTGTGAAAG
CAAAAAGCAATTGAGCCAGTTCGATCATGTACCGATTG{gtgagtcgaaactttgagacattca
agcttttccaataaacgtcttaatataaatttatataaatttaactgctggtttaattcataat
aacagcttatttcatagcatgggcggcgattagtgtcgtgatctgtttgtacttaacacttttc
taacagtttacgttttcacccgtcatag}GCATTCCACTTTCCAACACTGTTATCTACTTACTT
GATGGAGACTTCCGCCCAGTCAAATATGGCGAAATAGGTGAAATATTCGTTTCGGGCCTGAATC
TTGCCGAAGGTTATGTAAATGGTAGAGACCCGGAAAAGTTTGTCGAAAATCCATTGGCTGTGGA
ATTAA{gtaagttcaaacaaatcatccctctatgtaagtacgtaaatgaatacaatttctcttc
atcacaaacag}AATACTCACGTCTATATCGCACTGGTGATTATGGCTCTATACGCGATGGCAA
TGTGATGTTCGAAGGCCGCACCGATTCGCAAATAAAGATTCGTGGTCATCGTGTCGATCTTAGC
GAAGTTGAAAAGAACGTTTCAGAGTTGCCGTTCGTCGATAAAGCGATCGTACTTTGCTATCATG
CCGGCAAAATTGATCAGACGATTCTAGCTTTTGTTAAGCTACGCGACGATTCACCGCTAATCAA
CGAACTACAAATCGAGGCTAAATTGAGAGATAAGCTCGCGGAGTACATGTCGCCACAAGTTGTG
CTTATTGACCAAGTTCCATTCCTTGTCAATGGGAAAGTAGATCGGCAGGCACTTCTCAAAACTT
ACGAGACAGCCAATAATAACGAAGGCGATTCCAGTATTGTGCTGGACTATGATTATACACTCGT
ACCCGAAGAGGTAAAGCACATTGCCTGTGATCTCTTTGAGACCGTGGGTGGCGTAATCGGTCGT
TCTACACGCGCCACTCTATCGCCTCGCAGCAACTTTTACGAGTTGGGCGGAAATTCGTTGAATT
CGATTTACACAGTCACACTGTTGAGAGAAAAGGGCTACAATGTAGGCATATCTGAATTTATTGC
CGCAAAGGATTTGGGTGAAATATTGGAGAAAATGATAAATAATCGCAATGCCTCCCCAATGGTC
GACGACTGGACAAATGCTGCGCCACATCTTGATATGGTGGCTGAGCAGCTCAATGATTCCCACA
GGCAGCAAGTGGTGGA{gtaagtaaagcgatgaatgcattaattgcgtatgcatatctatatta

atttaattctctttttttacaaag}CATCATAGTTGATAGCTTTTATGGCAAGGCGGATCTGGAA
CAATGGCTCAAACCAGATATATTCCCAAATGACTACAGTGATCTTATTAAT{gtaatatttgat
ttaattccttccacttatgtatttataagtgaatttgttcttttccttgctatag}GATATGTG
GGATGTTTTAGTAGAGAAGAATCTCAGTTTAGTCGTGCGCGACAGGCGTACAAATCGCATCATT
GGAACGGCCCTAAATTTTGACGCACGCGCTGAACCCGAAGTCGAGGTTAAGTCAAAACTTATCA
TAGTATTTGAGTTTCTAGAGTTTGTTGAAGGCCCCATACG{gtaagattactacttatatatag
gcatatgtatatatactacataaacttatacacattcatacgcataagtacgtacatatgtaaa
gggtgatcaatttagaggtatcggattttaaattgaaataaaacaacgaaaattcaaattgatt
gggcaatctttattatttttgtatttatttttttaaagataatccctttcaaatgttggccgcaa
ctgcgccgtaattcggccatccgtaaacaccaattttgaatgaatcgcgactcgctggaggact
tcggtcagtatctggtatctagtgaataactttagtaatgttggcttccaatgcctcaattgaa
gctggtttatccacaaagcatttagactttacataccccagcatccccacaaataaaagtccga
aggtgtgatatcacacgatcttggtggccaatccactggtccgagacgagagataaattgctca
ccgaaacgacgacgcagtaaatccattgtttcacgggctgtatggcaagtagcgccgtcttgtt
ggaaccaaatattgtggagatcacgggcttcaatttcccaatttccggcatcaaaaagtcgttt
atcatggcgcgatagcgttcgccattcactgttacattggcgccagcctcgtctttgaagaaat
atgggccgatgattcctccagcccataggccgcaccaaacggttgttttcaatggatgtaatgg
ctgttcttgaatggcttcgggttgctcttcagcccaaatacggcaattttgcttattgacgtag
ccattaagccaaaaatgggcctcatcgctgaacaccataagttggcctgagcgcgcgatgaaca
cttttcacagagcgtcgattttcgtaatacaattgtacgatttgtaaacgttgttgaggcgtaa
gtctttccatgatgaaatgtcaatgaatactgaaaaaaattatgtatttagtttgacagtagtc
acgcgtgatctgtcaaaaaaaccctattggaaaaagtacctccaatctgatcaccctttatttc
accagatcacaaaaattttattatgtaacagtaaaaaggagacaagataaatatatacatccat
atacaaggtggcgcaaaattaatcaccctatcggaagatttataatttttgcaaatggcgtcgt
acgtcaatcatatttgacacttgtgaactagacagctgcagtatacaaacagacaagcaatgga
gcgcgtaaaggtcaaaataaagatttccactcaaaatcatttttttttttttttaaaaattttt
aagttatttaaaaattaaagtttataaaaagttattcaaaaaccaaatttgatgattaatttg
cgccacctttctctactcaagtaactaattcaatattgtcaagttagcatatttgaggcggcc
gccgtagccgaatgggttggtgcgtgactaccattcggaattcacagagagaacgtaggttcga
atctcggtgaaacaccaaaattaagaaaaacatttttctaatagcggtcgcccctcggcaggca
atggcaaacctccgagtgtatttctgccatgaaaaagctcctcataaaaatatctgccgttcgg
agtcggcttgaaactgtaggtccctccatttgtggaacaacatcaagacgcacaccacaaatag
gaggaggagctcggccaaacaccatgccatcaatcgagggaagctccaagtagctcattgcatg
cccgctaattgctattcatccaaaacacattacgactatatatcgaatgatcttaaaaatgaga
aattaaagaattatttaatggagggacaaaaacaaaaaaagttttatacaaattattcattttt
attttatttataatatttatccaaactaaaagtctatgcaaaactcaatttgcagttttataat
tgtttgcatttttttgctttaatttattttaaaataaag}GGACAACCAACTTCCCAAGGGGCTC
AACAAAATACTCCACTCATTCATGATGGGCACCAGCTCCGAATTGAATCCACAAGAGAATATCG
CCTGCATGTATTTCATGGAAAACGAAGTACTGCGCGTTGCTAGAGAGAAACAATTTGCCGGAAT
TTTAACAACCAATACAAGTCCCTTAACGCAG{gtaaaacaaaggaaccaaatgattccacgttt
tccaatatgagcgatttcttatacttttctccgtacaattacag}CAACTCGGTAACGATGTCT

ACCACTACAAGACATTGCTGGACTATCAGGTCAATCAATACGTATACATCGATGGCACTAAACC
ATTCGGAAAAGCTCCCGATTCACAACGTTCCATTGTTCATTGGCGCGAAGTTACCGACTAA

>Ceratitis_capitata_ebony
ATGGGCTCCATACCGAAATTGTCAATTGTAAAAGGCCAACAGCAGGACATAACGCCACGTCCTT
TGCATCGCATTTTCGAGGCCAACCTCCTACATCATTCACACAAAAGTGCGTTGATCACACACAA
AAATGCCGGCGGCACAATTACCTCCGAAGCCACACAATGCACCTATAAACAACTAAATCAATTG
GCCAACCAATGTGCGCGCCTACTGGTGAGCGACATAAAACGTAATTGCTTGGAGCCGAATACAG
ATGGGGATTACATTGTGGCGGTTTGCATGGAACCATCAGATGCACTTGTCAATACGCTGCTCTC
CATTTGGAAAGCCGGTGCCGCTTATCTGCCAATAGATCCCACTTTTCCGCTCAATCGCGTACAA
CACATACTCCAGGAAGCGCGTCCCGTTATGGTGCTATGTGATGACAATATTGAGCGTGAACGTT
TCAACGATACACCGACCATCAGCAGCTCGGAACTCTTTCATCGTTCTGCCGATTTGAGTAATGC
TAATTTATTACCAGCTGAAATGCTTGCTGACGGTGCAAATAATTTGGCCATTGTGCTATACACC
TCTGGGAGCACCGGTGTGCCAAAGGGTGTGCGTTTACCGCATGAGGTCATCTTAAATCGTTTGC
AATGGCAATGGCATACATTCCCTTATGCGTCCACAGAGCAGGTGAGCGTTTTTAAAACTGCGCT
AACATTTGTCGACTCCGTCTCCGAGCTGTGGGGCCCACTTATGTGCGGTCTTTCGATTTTGGTT
GTACCAAAAGTGATAACTAAAGATCCCGAACGTTTGGTGGATCTATTGGAAAAATATAAAATAC
GTCGTTTGGTTTTGGTGCCAACACTTTTACGTTCGATTTTAATGTATTTAAAGTTGAATGATGA
CAATACCAAGCGTCAGCTGTTACAACGCGATCGCCTGTTGTATAATATGAAGATTTGGGTTTGT
TCCGGTGAACCGTTGCCAGTAGCTTTGGCTACGAGCTTCTACGATTACTTTGCTGAGGGCGTAC
ATACATTATTTAACTTTTACGGTTCCACCGAAGTGATGGGCGATGTCACTTATTTTGCTTGTGA
AAGTAAGAAGCAGTTGAGCTTCTTTGATCATGTACCAATTG{gtaagtaaatattttgtttgtt
tgtttttaagttcaagctgcagtctatggtttcgcggtctgaagaagagttatgtgaggtaaa
acatgagtcaagataggtagagcgtaatctgctacaaatatataattgcgccattactgtttct
tgagctcttcatttgcattaattggtggttagattctgataacgtcattgggtgaggacataga
tgaaattgtgaagtcattgcggccggacgtgttttgtctacgctcgagagtttaatactgtttc
ggtggcattttaaataaaaattcttggtattgtttcagtgagacagtttgtttgtttgtctacc
cgttgaagctgatttcttgtatatctttttgtatttgtgcttatattcgtcagtttactttt
aaataaatatgagcacctcagccagccaatccagcctaaaacgccgtgctagcagagaatcacc
gttcgccaagtctcttaagcttgacgatcacaataataatgctctccccactttactgactgca
gtacgagatttggaggagagagtgaaaatgcaaactgaagagatgcactcactttttaacacgc
tctctggcaaaatcaatgatatgcaagagatgctcatgaagaagctgagtgatgaagtcagcgc
attaaatcttaagtgtaatgagtctgctgagcgcgttttgcgacttgaggctgatatgaaaaaa
caagaaggtgaaatgcagaaacttaaagcaacaataaacttgctgtcaagcagaggtgagagtt
ctcaggcagatgcaattgcttttggtgtaccatttacggagggagagaacattaggagtatctt
caaccaagtatgtctctccatcgactttatggtaccacagatacgtgatgcgttcagaataaaa
cagtcgaatatgataattataaaattttattctcctgctgatcgcgcgcgtactcttcgcgcat
tttctgaatgtcgaaagagaataaaatcggctattcccctaacatctataggaattgatggcaa
gtttagcatcttcgagagcctgtctaccgagaaaagaaaaattttacaagaaggtttaaagcta
aaacgtttggggaagctgcagtcggtgttctctatgagaggagatgtttatgttagactcagta
cagactccactgcatgcaaaataggaaatgttgatgatcttcagcgatttcgttaacattaact

tgttatatttatatttatatttttatttgtattgtagctttatatatatttcatatatgtttta
tttgaatgtttttgtaaaaatggtcattctcttttccttgctttatattgttattcttgaggtt
gcttatgtttttgttcttgttgagcatgcaaaacagctgtcctagtattggcatgaacagtctc
atgattgatgtgctatgtaaacaaaacagtggattcaacatagtccatttcaatgcgcggagtc
taagttcagataaaatggattatgttcgcaacacatttgagctttcatcaattgatgtcatctg
tgtttctgaaacatggtttgatgaggatttatgttactcgtctactaacataaaaaactacaaa
ttaatacgtcatgacaggccaggaagaagaggaggtggggttgcaatattttgcaaggacatat
tttctatgaaattactttcaaaatctgctgatagcgatattgaatatttaattgtagaagtaag
ccacaaaaaattccagactgttgtttcttgtgtttataatcctcaccgaactaatgttgtcaaa
ccttttttgatgtaatttctttatatgttgttaagtacaattcagttatagtatgtggtgatt
ttaatgttaatcttcttgtcaacgactcttatacaaataatttacttgacaatctatccagttg
ctctctctctgtggtcaaccgatctcttcctacgcggtttgcagtgaattgcactcctagtctt
cttgactacttcataattcttgacttaactactgtattaaaatttgaccaattaacttttatat
ctgaccatgatctcattttctgctgtttggacataaatatcgatcgttgttgcataaataagac
tatctcttttagagattaccgctttgtcaacacaaatttacttctttcggaattgtattgtgta
gactggagtgaatgctggatatatccatctaccgatgaaaaacttgatttcttaatgggaattt
taaagtcagccttcgagagacatgtccccattcgtacaatcactaataagtctatgtcctgtcc
atggtttacctcaaaatctctgaaatcaattaagttacgaaataaacttcattccatatggaaa
aagaaacccacgttggaaaattggaatgcctttaaatcggctcgaaacagagcgacgcttgtta
tcagaaatgaaaaacgagcttattttaacaagaaactaaattctggcctctcaacaaaggttct
atggcgtaacattaaacgacttggtgttcattgtaaggagaatgtcgaatgttccttagaagca
aatgtggttaatgatgcttttcttctaaactgtgtaccacacagtgacactattaactttatgt
ccagggatgatctcttgtttagggatgactgttttctattttcagctgttagtgaatatgatgt
tgcaatgagtatatttaaaatctcatccaatgcaatagggcatgatggtttaccaattaaattt
attaaaattatactaccttgcattcttgggacactaacgcacattattaatcattgtattacca
cgtcttgtttcccagagcaatggaaaattggcactgtactacctgtcgctaaaaagaaccgtcc
ttgtagtccgtctgattttcgaccaataagcgtcctccctgtattgtctaaagtttttgagatg
cttttagctaagcaaataaacgaatttgtaagcaagcataacttaatatctccgtttcaatcag
gcttcaggaaaaatcacagctgtagtactgcggtactaaagatcagtgatgatattcgatccaa
catggatgaaaataaattcaccttgctttgcctacttaacttttcaaaagcatttgatatggtc
aaccatgagttattaagctgtaagttgaaagcgtactttggttttagttgtagtgctttaaaac
ttgtgaaaagctaccttgttgacaggtttcaatttgtaaaaattggggaggactcatcgcaact
caagccagttacttcaggggtgcctcaaggctccatccttggtcctctgcttttcagcatattt
attaatgacatcttcagtgtgtgcaaattttcaaacctgcatggttatgctgatgacattcagt
tatatttatctgcaccttttgatcaagctaaagagttatgtaaaagcattaatactgacttggc
tgatatctcttgctgggcggctgccagcggtctaattttaaataaaaacaagtgctttgtgctg
ccaatcagtgctagaaaactgacaaatctggatttttttgcccccaatattcatagacaattatc
ccttaaaattagtcagcaagattaaaaacttaggatacattattaactctaatcttacatgcac
ggatcatgtaaactccatagttagtaaggtctatctcattctgaggaatcttaggcagtcagct
atctatacacccactgctaccagacgccgacttgctatacaaattattcttccggtcatcatgt
actctgaagtcgtatacagcaaacatgacagtttttctgctcgtaaaattgacgttatgtttaa

taatgtcacacggtacgtgtttggtctttccaaatatgagcacatttcgtcttggaaagatcag
atattaggatgtacgattgcggactatcataaagccagaaattgtatttttctttgcaagctca
tcatgactcaatcacctagttatctttttccaagttatcgtttactcggtcagccagagtagg
taatataataatacctttcatagatcacagcaatcatcaaggctgttctttgtcaatgctagt
cgattatggaactctcttcctgctaatttaaggaggaaattaaacagaagcaatttcaagaacc
acatatatatgcattttaaaggccaaatttcaaacatagactgaccattcaataattgtttaag
tttttgtcatactcaactcattatatattataattatttttatttttattattatttttgttaa
ggtatctgtaaagtaatatttacttattcttcacactatctctgttgtactaaaaaagatatgg
aaatcttattgtactaatgctttgttattattattaaataaataaataaataaataaataaata
gaatgttctgtctaagttcgtataagcaacaagagctggtggtgctctaactggtctccaactt
tttcatcctttagtcaccagtctcgatcagcaaaagaacaaaagtatttcccttattaacatta
taatgaactttaacaattccaatgttttaagggtttgcaattcgaatagggtttgaacgacgac
atcaatactaagagaaaataagacaatatttaccaaaactatatggagagcgtggtattacgac
gtccttcccgatgtactttggaaactgttgtgctttgagcaagtgtctaatcctaaccagagta
atggtttcgtattgaatggtgaagactcgataattcactagttacgtatgcggaatcttatgag
aagcgactgtagtaagggcctttgaattctcattgagagaggtactttagtaaatagttagtgg
atctaataatctcaaaataaaaaacccttgataggctttttatcctcaattaaatacttgaatg
tatcttttttgagacttttgatgcaaagaattatttttttcagtaattttcaatttcgtttagcc
taggctggcgtgatttgttgattcagttcctcagttccatcatttctttttcatataatttgag
tagttgatttctaattaactataaagataaactcttaaatatgtagcatttccactcttccag}
GCATACCACTTTCCAACACTGTCATCTATTTACTCGACAGCGACTTCCGACCAGTCAAACAAGG
AGAAATCGGTGAAATCTTCGTTTCAGGACTCAATTTAGCTGAAGGTTATGTAAACGGCAGAGAT
CCAGAAAAATTTGTAGAAAATCCATTAGCCGTGGAGTTTA{gtaagttcaaacaaataattttc
tgataaaatgtacaatacgtaaattgtaatccccactttcatcaccaaag}AATACTCCCGACT
CTATCGCACTGGCGATTATGGATCTCTACGTGATGGCAATGTTATGTACGAAGGACGCACTGAT
TCACAGATTAAAATACGTGGTCACCGCGTTGATCTTGCGGAAGTTGAGAAGAATGTTTCGGAAT
TGCCGTTTGTCGATAAGGCAATGGTATTATGTTATCATGCTGGAAAAATTGATCAAACGATTTT
GGCATTTGTCAAACTACGCGACGATTCGCCACTGATAAGCGAGTTGCAAATCGAAGCGAAATTA
AGAAGCAAACTTGCTGAGTATATGACACCACAGGTTGTTTTAATCGATAAAGTGCCTTACCTTG
TCAATGGGAAAGTGGATCGGCAGGCGCTACTCAAAATGTATGAAACAGCAAATAACAACGAGGG
TGACTCGAGCATTGTACTCGACTACGATTACTCACTGGTGCCCGCGGAGATTAAGCACATAGCT
GTGGATCTCTTTGAGACCGTGGGCGGTGTGATTGGACGTTCTACACGCACTACTTTGACGCTAC
GCAGCAACTTTTACGAATTGGGTGGCAATTCGTTGAATTCGATTTATACCGTGACTTTGTTGCG
TGAAAAAGGCTATAATGTGGGTATATCTGAATTTATTGCGGCAAAAGATTTAGGTGAAATATTG
GAGAAAATGGTGAATAACCGTAATGCGACGACAATGGTCGAAGACTGGAGAGGTGCTGCACCAC
ACCTAGATATGACCGCTGAACCGCTCAGTGATGTACACAGGCGAGAAGTGATTGA{gtaagttt
gacgatcaaaatgtatgtaaaaaaaatttaaaaacaaaactctctttcttctttaaag}TATCA
TAGTTGACAGCTTCTATGGCAAAGCGGATCTTGAACAGTGGTTGAAGCCCAATATATATCCAAA
TGATTACAGTGATTTGATAAGT{gtaagtttctaatatcactgaatttccttactctgcttgca
cagaagagtgcacttacatatagtatgtacatatataatttctatcttcattctcctcgttcta
tag}GACATTTGGAACGCTTTGGTTGAGAAGAATCTCAGCCTGATTGTACGAGATAAGCGCTCA

AATCGTATAATTGGTACGGCATTGAATTTCGATGCTCGCGCTGAGCCAGAAGTTGAGGTTAAGT
CGAAATTAATTATCGTTTTCGAGTTTTTGGAATTTATTGAAGGACCTATAAG{gtgagtttctt
acaataaattattaataattgatatttatgtgtgtaggtatattcgctgcgctatgttattatt
ctgcggctttgggttgcaatctatacttatacggtcaagcatagatccgattgtcattcacgcg
acgggctttcggtagcgaggaatataatacaaaaaaaaacttattattgggtgtatgcattaaa
tcgtggggttttctaagagagggctctactagtaatattatgcgtcgtgttcatctgtggctat
attcatttcaaagggactgtcatttccgaacgaacatgccaaaagtggtttgcacgctttaaaa
gtggcaatttcgaattagacgacggatagccaaaaaaatttcaagacgaagaattagagacatt
gctcgacgcagatacatgccagacgcaagaagaacttgcaaaagtattaggagttgatcaagca
accgtttccagacgattaaaatcaattttacaaaaaagtcttaattttcgggggaaaaaactc
atgatttagtgcatacaatcatatttgttattgtatgatgtttgccaaggggttttggtcatta
tttgcaatattaatgagagcaaaattctacaaaatacataaagccacaagaacctgtattttag
tatatacttgcatgcatatatatatgtgttttcttatttttcaatgcttttatatttgattttt
tttaatgctttctttttcgcaatatag}AGACAACCAACTACCCAAGGGCCTCAACAAAATACT
GCACTCATTTATGATGGGCACCAGTTCGGATTTGAATCCACAAGAGAATATTGCCTGCATGCAT
TTCATGGAAAACGAAGTGTTACGCGTTGCCAGGGGGAAAAAGTTTGCCGGCATTTTAACCACTA
ATACAAGTCCCTTGACACAG{gtgaaataatatacatatatattttttagttttcaattaatt
tattttacaatttttcacttatccttttag}CAATTGGGCAACGATGTCTACAACTACAAAACC
TTATTGGACTATCAAGTCAATCAGTATGTGTATAACGATGGCACGAGACCCTTTGGAAAGGCGC
CTGATTCACAACGCGCCATCGTTCACTGGCGTGAGGTGACTGACTGA

>Bactrocera_tryoni_ebony
ATGGGATCCATACCGAAATTGTCAATTGTGAAAGGCCAGCAACAGGACATTACACCGCGTCCAT
TGCATCGCATTTTCGAAGCCAATCTGCTGCATCACTCACACAAGAACGCACTAATAACACACAA
CAATGGGGACTCCATTGCCTCCGAGTCCACACAATGCACCTACAAGCAGTTAAATCAGCGCGCT
AACCAATGTGCACGTCTGCTAGTAGCTGATATAAAACGTAATTGCCTGCAGCCTAACACCGATG
GTGATTACATTGTGGCGGTATGCATGGAGCCGTCCGATGTGCTTGTCAATACACTGCTCTCGAT
TTGGAAAGCCGGTGCTTCTTATCTACCAATAGATCCAACCTTTCCGAACAATCGTATACAACAC
ATATTGCAGGAATCACGTCCCGTTATAGTGCTATGCGATGACAATATCGACCGTCAACGCTTTA
ACGGTACACCGACCATCAGCAGTTCTGAGCTTTTCCATCGTTCGGCCGAATTGAGTAACGCTAA
TTTAATGCCGGCCGAAATGCTAGCCGACGGTACTAATAATTTGGCAATTGTGCTTTACACATCG
GGCAGCACTGGCGTACCCAAGGGTGTGCGTCTACCACACGAGATAATCTTGAACCGTTTGCAAT
GGCAATGGCATACATTTCCCTATGCGCCGACTGAGTCCGTGAGCGTTTTCAAGACCGCACTCAC
CTTTGTTGATTCGGTGTCGGAATTGTGGGGACCGCTGATGTGTGGTCTTTCTATTTTGGTTGTG
CCAAAGTCGATAACTAAAGATCCGGAGCGTTTGGTGGATTTGCTGGAGAAATACAAAATACGTC
GTTTAGTTTTGGTGCCGACACTTTTACGTTCGATTTTGATGTTTTTGAAGTTGAATGATGGCAA
TACTAAATGTCAAATGTTGCAGCGCGGTCGTCTACTGTATAATTTGAAGATTTGGGTTTGTTCC
GGTGAACCATTACCAGTAACTTTGGCTGCGAGTTTCTTCGACTATTTTTCCGAAGGCATACATA
CGCTTTACAATTTTTATGGATCCACCGAAGTGATGGGCGATGTCACTTATTTTGCTTGTGAAAG
CAAAAAGCAATTAAGCTACTTCGATCATGTGCCGATTG{gtaagtgaattttctcaccgagctc
taataatgatatttaaaaagcacttctaatgactcatttatgtatgtaattatatcttcacag}

GTATTCCACTTTCAAACACTGTGATATATTTACTTGACGGCGATTTCCGACCAGTCAAACAAGG
AGAAATTGGTGAAATTTTCGTTTCGGGTCTAAATCTAGCCGAAGGTTATGTAAACGGTAGAGAT
CCAGAAAAGTTCGTGGAAAACCCCTTGGCTGTGGAATTTA{gtaagatcaacgtaactcaaaca
cgtataaaaataccaatatatcttttctttacgcag}AATACTCACGCCTCTATCGCACCGGTG
ATTACGGCTCTTTGCGTGACGGCAATGTAATGTACGAAGGACGCACTGACTCGCAAATAAAAAT
TCGTGGTCATCGTGTTGATCTTGCCGAAGTTGAGAAGAACGTTTCAGAATTACCATTCGTCGAT
AAAGCGATCGTATTGTGCTATCATGCTGGCAAAATTGATCAGACAATTCTGGCTTTCGTAAAAT
TACGCGACGATTCACCACTACTGAGCGAAATGCAAATTGAAGCGAAATTGAGGGACAAACTTGC
TGAGTATATGACACCACAAGTGGTTCTAATCGACAAGGTGCCCTACCTTGTTAACGGTAAAGTA
GATCGGCAGGCACTACTAAAAACTTACGAGACGGCAAACAATAACGAAGGTGACTCCAGCATTG
TACTGGACTACGACTATACGCACGTGCCCGAAGATGTCCAGCATATAGCCAGGGATCTTTTTGA
GACTGTGGGTGGTGTGATTGGGCGCTCAACACGCACTACTCTATCGCTGCGCAGCAACTTTTAT
GAGTTGGGCGGAAACTCATTGAATTCCATTTATACAGTTACGTTGCTGCGTGAGAAGGGCTACA
ATGTTGGCATATCCGAGTTTATTGCGGCTAAAGATTTGGGTGAAATATTGGAGAAAATGGCACA
TAACCGCAATTCGAATGCAATGATCGAGGACTGGTCGAATGCTGCACCACATCTCAACATGATC
GCTGAACCGCTTAGCCATGCGCATAAGCAGGATGTAGTTGA{gtaagtaatgcgaaattgttgg
tcaagatcgagtgtgcttaaatgcttttgtctattattttacatag}AATCATCGTCGACAGCT
TCTATGGTAAGGCGGATCTAGAGCAATGGCTAAAACCTGACATTTTTCCAAACGATTACAGTGA
TCTAATCAAT{gtaagaatttcatcggccttcattggccgcttcttcacttcgttaagattgtt
taataatttatctatttaatag}GATATATGGGACGCATTGGTGGAAAAGAACCTAAGTTTGGT
GGTGCGTGATAAGCGCACGGATCGTATCATTGGTACTGCGTTAAATTTTGACGCACGCGCTGAG
CCAGAAGTTGAGGTTAAATCGAAATTGATTATTGTTTTCGAGTTTTTGGAATTCGTTGAAGGTC
CTATACG{gtaagtcctatgcggtaatgggtaaaccaatatgaaagcgcagtaaaaagcagctg
atctttttagaaaaaataataattttcactccttacttttgaaatcctaaagttcgtaaatgtt
gatattgatgctcgatttttataaatatattttgaaaaatttattttttcgaaacaaag}AGA
CAATCAACTGCCGAAAGGACTCAACCAAATACTGCACTCATTCATGATGGGCACCAGCTCCGAT
TTAAATCCACAAGAGAATATTGCCTGCATGCATTTCATGGAAAATGAGGTGCTTCGCGTTGCTA
GAGAGAAGCACTTCGCTGGCATCTTAACAACCAATACGAGTCCCTTAACGCAG{gtaagttaag
ttagttaatagaaaccttccttgttttcagtacaattccaatataagttacattaacggtctgc
atttctttcttcttcactccttcag}CAATTGGGCAACGATGTCTATCAGTATAAAACATTGCT
GGACTACCAGGTCAATCAATATGTTTACAACGATGGTACAATACCCTTTGGCAAAGCGCCCGAT
TCGCAACGTGCCATTGTCCATTGGCGCGAGGTGACCGACTGA

>Bactrocera_dorsalis_ebony
ATGGGATCTATACCGAAATTGTCAATTGTGAAAGGCCAACAACAGGACATTACACCGCGTCCAT
TGCATCGCATTTTCGAAGCCAATCTGCTGCATCACTCACACAAGACCGCGCTAATAACACACAA
CAATAGCGACTCCATTGCCTCCGAGTCCACACATTGCACCTACAAGCAGTTAAATAAGCGCGCT
AACCAATGTGCACGTCTGCTAGTAGCTGATATAAAACGTAATTGCCTGCAGCCCAACAACGATG
GTGACTACATTGTGGCGGTATGCATGGAGCCGTCCGATGTGCTTGTCAATACACTGCTCTCGAT
TTGGAAAGCCGGTGCTGCTTATCTACCAATAGATCCAACCTTTCCGAACAATCGCATACAACAC
ATATTGCAGGAATCACGTCCCGTTATAGTGCTTTGCGATGACAATATCGACCGTCAACGCTTTA

ACGGTACACCGACCATCAGCAGCTCTGAGCTTTTCCGTCGTTCGGCCGAATTGAGTAACGCTAA
TTTGATGCCGGTCGAAATGCTGGCCGACGGTACTAATAATTTGGCAATTGTGCTTTACACATCG
GGCAGCACTGGCGTACCGAAGGGTGTGCGTCTGCCGCACGAGATAATCTTGAATCGTTTGCAAT
GGCAATGGCATACATTTCCCTATGCGCCAACTGAGTCCGTGAGCGTTTTCAAGACCGCACTCAC
CTTTGTTGATTCGATATCGGAATTGTGGGGACCGCTGATGTGTGGTCTTTCTATTTTGGTTGTT
CCAAAATCGATAACTAAAGATCCGGAGCGTTTGGTGGATTTGCTGGAGAAATACAAAATACGTC
GTTTAGTTTTGGTGCCGACACTTTTACGTTCGATTTTGATGTTTTTGAAGTTGAATGATGGCAA
TAATAAATGCCAAATGTTGCAGCGCGATCGTCTACTGTACAATTTGAAGATTTGGGTTTGTTCC
GGTGAACCATTACCAGTAGCTTTGGCTGCGAGTTTCTTCGACTATTTTTCCGAAGGCATACATA
CGCTATACAATTTTTATGGATCCACCGAAGTGATGGGCGATGTCACTTATTTTGCTTGTGAAAG
CAAGAAGCAATTAAGCTACTTCGATCATGTGCCGATTG{gtaagtgaattttctcaccgagctc
tattaataatatttataaagcacttctaataactcacatatgtatgtacttatatcctcacag}
GTATTCCACTATCAAACACTGTGATATATTTACTTGACGGCGATTTCCGACCAGTCAAGCAAGG
AGAAATTGGAGAAATTTTCGTTTCGGGTCTAAATCTAGCCGAAGGTTATGTAAACGGTAGAGAT
CCAGAAAAGTTCGTGGAAAACCCCTTGGCTGTGGAATTTA{gtaagtccaacgtaactcaagca
cgtataaaaataccaatatatatatttctttacgcag}AATACTCACGCCTCTATCGCACCGG
TGATTACGGCTCCTTACGTGACGGCAATGTAATGTTCGAAGGACGCACTGACTCCCAAATAAAA
ATTCGTGGTCATCGTGTTGATCTTGCCGAAGTTGAGAAGAACGTTTCAGAATTACCATTCGTCG
ATAAAGCGATCGTATTGTGCTATCATGCTGGCAAAATTGATCAAGCAATTCTGGCTTTCGTAAA
ATTACGCGACGATTCACCACTACTGAGCGAAATGCAAATTGAAGCGAAATTGAGGGACAAACTT
GCTGAGTATATGACACCACAAGTGGTTCTAATCGACAAGGTGCCCTACCTTGTCAACGGTAAAG
TAGATCGGCAAGCACTACTAAAAACTTACGAGACGGCAAATAATAACGAAGGTGACTCCAGCAT
TGTACTGGACTACGACTATACGCACGTACCCGAAGATGTCCAGCATATAGCCAGGGATCTTTTT
GAGACTGTGGGTGGGGTGATTGGGCGCTCAACACGCACTACTCTATCGCTGCGCAGCAACTTTT
ATGAGTTGGGCGGAAACTCATTGAATTCCATTTATACAGTTACGTTGCTGCGTGAAAAGGGCTA
CAATGTTGGCATATCCGAATTTATTGCGGCTAAAGATTTGGGTGAAATATTGGAGAAAATGGTA
CATAACCGCAATTCGAATGCAATGATCGAGGACTGGTCGAATGCTGCACCACATCTCAACATGA
TCGCTGAACCGCTTAGCCATGCGCATAAACAGGATGTAGTTGA{gtaagtaatgcgaaactgat
ggcgtcaagttcaaatgtgctaaggtgcttttgtctactcttttacatag}AATCATCGTCGAC
AGCTTCTATGGTAAAGCGGATCTGGAGCAATGGCTAAAACCTGACATTTTTCCAAATGATTACA
GTGATCTAATCAAT{gtaagaatttcatgccctcattggctgcttcttcactttgttaagattg
ttcaataatttatctgtttattag}GATATATGGGACGCTTTGGTGGAAAAGAATCTAAGTTTG
GTGGTGCGTGATAAACGCACGGATCGTATCATTGGTACGGCATTGAATTTCGATGCACGCGCTG
AGCCAGAGGTTGAGGTGAAATCGAAATTGATTATCGTTTTTGAGTTTTTGGAATTCGTTGAAGG
TCCTATACG{gtaagtcatatgcggtaattgttaaatatataggaaagcgcagtaaaaagagct
ggccttttttagaaaaagaaaataataatttccactgcctacttttgaatttctgaaatttgtga
atgttgatattcacgttcaattttatatatacttacatacaagtacatatgtgcatacctatat
ttcgaatatttttaaaatttaatttttctaaacaaag}AGACAATCAACTGCCGAAAGGACTC
AACCAAATACTGCACTCATTCATGATGGGCACCAGCTCCGATTTAAATCCACAAGAGAATATTG
CCTGCATGCATTTCATGGAAAATGAAGTGCTGCGCGTTGCTAGAGAGAAGCACTTCGCTGGCAT
TTTAACAACCAATACGAGTCCCTTAACGCAG{gtaagataaattagttaatgaaagtcttcctt

gttttcaatacgattccaatataaattatatttaacgatgtgcatttgtttcactcactcttgc
ag}CAATTGGGCAATGATGTCTATCAGTACAAAACATTGCTGGACTACCAGGTCAATCAATATG
TGTACAACGATGGCACTGTACCCTTTGGCAAAGCGCCCGATTCGCAACGTGCCATTGTCCATTG
GCGCGAAGTGACCGACTGA

>Zeugodacus_cucurbitae_ebony
ATGGGCTCAATACCGAAATTGTCAATTGTGAAGGGCCAACAGCAGGACATTACAGCACGTCCAT
TGCATCGCATTTTCGAAGCCAATCTGCTGCATCACTCGCACAAAAGCGCACTCATAACACACAA
CAATGGCAGCGCCATTGCCTCCGAGTCCACACAATGCACCTACAAGCAGCTAAATCAGCGCGCC
AATCAATACGCACGTCTTCTCGTGGCCGATATAAAACGCAATTGTTTGGAACCCAATAACGATG
GTGACTACATTGTGGCCGTATGCATGGAGCCATCCGATGTGCTGGTCAATACACTGCTCTCCAT
TTGGAAAGCTGGCGCTGCTTATCTACCAATCGATCCCACTTTCCCCATTAATCGCATACAACAT
ATTCTGCAGGAAGCACGTCCCGTTATGGTGTTGTGTGACGACAATATTGAGCGACAACGTTTCA
ACGGCACACCGACACTCAGCAGCTCTGAGCTTTTCCAACGTTCGGCCGAATTGAGTAACGCCAA
TTTAATGCCGGCCGAGATGTTGGCCGATGGCAGCAACCATTTGGCCATTGTTCTCTACACATCT
GGCAGCACTGGTGTACCGAAAGGTGTGCGGTTACCGCACGAGGTGATCTTGAATCGTTTGCAAT
GGCAATGGCATACATTCCCCTATGCGCCGACCGAGTCCGTGAGCGTTTTCAAAACCGCGCTCAC
CTTTGTGGACTCGGTATCGGAATTGTGGGGGCCGCTAATGTGTGGTCTCTCCATTTTGGTTGTG
CCAAAGTCGATAACTAAAGATCCCGAACGTTTGGTGGATTTATTGGAGAAATATAAGATACGCC
GTTTAGTTTTGGTGCCGACACTTTTACGTTCGATTTTGATGTTTTTGAAGTTGAacgatggcaa
caacaaataccaaaTGTTCCAAAGTGACAGACTCTTGTATAATTTGAAGATTTGGGTTTGCTCT
GGTGAACCATTACCGGTGGCATTGGCTTCGAGTTTCTTCGACTATTTTTCCGAGGGTGTACATA
CGCTCTACAATTTTTATGGTTCCACCGAAGTAATGGGTGACGTCACATATTTCGCTTGTGAAAG
CAAGAAACAATTGAGCCTCTTCGATCATGTGCCGATTG{gtaagtcaactactccgctgaggtc
tgggtattataaattgtactctctcatatttaaatccccacag}GCATTCCACTTTCCAACACT
GTAATCTATTTACTCGACGGCGATTTCCGTCCAGTCAAACAAGGTGAAATTGGTGAGATTTTCG
TTTCCGGTCTCAATCTAGCCGAAGGTTATGTCAACGGTAGAGATCCAGAGAAGTTTGTAGAAAA
CCCTTTGGCTGTGGAATTTA{gtaagtacaaagtcactcatacaaaatataaaatatttcaatc
ctcctctcctccctctcag}AATACTCACGTCTCTATCGCACTGGTGATTATGGTTCCCTACGC
GACGGCAATGTGATGTACGAAGGACGCACCgattcacaaataaaaattcgTGGACATCGTGTTG
ATCTAGCCGAAGTTGAGAAGAACGTTACCGAATTACCGTTCGTCGATAAGGCGATGGTCTTGTG
TTATCATGCTGGAAAAATTGATCAAGCCATACTGGCTTTCGTAAAAATACGTGACGACTCACCG
CTACTCAACGAAATGCAAATTGAAGCGAAATTGCGGGACAAACTCGCCGACTATATGACACCAC
AAGTGGTGCTCATCGATAAGATGCCCTATCTCGTCAACGGCAAAGTAGACCGACAGGCACTACT
AAAATCTTATGAGACCGCAAATAATAACGAGGGTGACTGCAGCATTGTGCTGGACTACGATTAT
ACACTCGTACCCGAAGAGCTCAAACACATCGCCAGGGATCTCTTCGAGACCGTCGGTGGCGTAA
TCGGGCGCTCAACACGCACTACGCTTTCGCCACGCAGCAACTTTTATGAGTTGGGTGGCAATTC
ATTGAATTCGATTTATACAGTTACTTTGCTGCGCGAGAAGGGTTACAATGTGGGCATATCTGAA
TTTATTGCGGCCAAAGATTTGGGTGAAATACTGGAGAAAATGGTAAATAATCGCAATTCGAATG
CAATGATCGAGGACTGGTCGAATGCTGCACCACATCTAGACATGATCGCCGAACCGCTGAGCGA
TGCACACCGACAAGATGTGGTTGA{gtaagtattacgaatttttttaggttacagatatatattt

aagtgtcgtcttcacttttacacag}TATCATAGTGGACAGCTTCTATGGCAAGGCGGATCTGG
AGCAATGGCTGAAACCCGATATTTTCCCTAGTGACTACAGCGATCTAATTAAT{gtaagacttt
gatttcattatttgtgaaggcctctaacctaaattgttacttggattacattcaatttttctat
tctatag}GAAATATGGCACGCTTTGGTCGAGAAGAATCTTAGTTTGGTGGTGCGTGATAAACG
CACGAATCGCATCATTGGTACGGCCTTAAATTTTGACGCACGCGCCGAACCAGAAGTTGAGGTT
AAGTCGAAATTGATAATTGTTTTCGAGTTCTTGGAGTTCGTTGAAGGTCCCATACG{gtaaata
gcaagcacatgaagcgataaagttacaataaataatttttctttattattattctaaacaaaag
}GGACAATCAACTGCCGAAGGGACTAAATCAGATACTACACTCATTCATGATGGGCACCAGCTC
CGATATGAATCCGCAAGAGAATATCGCCTGCATGCATTTCATGGAAAACGAAGTGCTACGCGTT
GCCAGAGAGAAGCAATACGCCGGAATTCTAACAACCAACACGAGTCCGCTAACGCAG{gtgagc
taactaacataaaatactccactgatctctatattatatattattttccacttgtatattcag}
CAATTGGGCAACGATGTCTATCAATACAAAACATTGCTGGACTACCAGGTCAACCAGTATGTAT
ACAACGACGGCACACAACCCTTTGGTAAAGCACCCGATTCACAACGTGCTATTGTCCATTGGCG
CGAAGTGACCGACTGA
